# Ligand-Dependent Mechanisms of CC Chemokine Receptor 5 (CCR5) Trafficking Revealed by APEX2 Proximity Labeling Proteomics

**DOI:** 10.1101/2023.11.01.565224

**Authors:** Siyi Gu, Svetlana Maurya, Alexis Lona, Leire Borrega-Roman, Catherina Salanga, David J. Gonzalez, Irina Kufareva, Tracy M. Handel

## Abstract

CC chemokine receptor 5 (CCR5) promotes inflammatory responses by driving cell migration and scavenging chemokine to shape directional chemokine gradients. A CCR5 inhibitor has been approved for blocking HIV entry into cells. However, targeting CCR5 for the treatment of other diseases has had limited success, likely because of the complexity of CCR5 pharmacology and biology. CCR5 is activated by natural and engineered chemokines that elicit distinct signaling and trafficking responses, including receptor sequestration inside the cell. Intracellular sequestration may be therapeutically exploitable as a strategy for receptor inhibition, but the mechanisms by which different ligands promote receptor retention in the cell versus presence on the cell membrane are poorly understood. We employed live cell ascorbic acid peroxidase (APEX2) proximity labeling and quantitative mass spectrometry proteomics for unbiased discovery of temporally resolved protein neighborhoods of CCR5 following stimulation with its endogenous agonist, CCL5, and two CCL5 variants that promote intracellular retention of the receptor. Along with targeted pharmacological assays, the data reveal distinct ligand-dependent CCR5 trafficking patterns with temporal and spatial resolution. All three chemokines internalize CCR5 via β-arrestin-dependent, clathrin-mediated endocytosis but to different extents, with different kinetics and varying dependencies on GPCR kinase subtypes. The agonists differ in their ability to target the receptor to lysosomes for degradation, as well as to the Golgi compartment and the trans-Golgi network, and these trafficking patterns translate into distinct levels of ligand scavenging. The results provide insight into the cellular mechanisms behind CCR5 intracellular sequestration and suggest how trafficking can be exploited for the development of functional antagonists of CCR5.

**Significance Statement:** CCR5 plays a crucial role in the immune system and is important in numerous physiological and pathological processes such as inflammation, cancer and transmission of HIV. It responds to different ligands with distinct signaling and trafficking behaviors; notably some ligands induce retention of the receptor inside the cell. Using time-resolved proximity labeling proteomics and targeted pharmacological experiments, this study reveals the cellular basis for receptor sequestration that can be exploited as a therapeutic strategy for inhibiting CCR5 function.

## Introduction

CC chemokine receptor 5 (CCR5) is a G protein-coupled receptor (GPCR) that mediates migration and activation of immune cells in response to chemokines CCL3, CCL4 and CCL5 (1). Because of its role in facilitating entry of the human immunodeficiency virus (HIV) into leukocytes during viral transmission, CCR5 was intensively studied as a therapeutic target for the prevention and treatment of HIV (2, 3), resulting in FDA approval of the small molecule CCR5 antagonist Selzentry^™^ (Maraviroc) (4). It has also been investigated because of its role in inflammation associated with Alzheimer’s disease (5), nonalcoholic steatohepatitis (6), multiple sclerosis (7), atherosclerosis (8), inflammatory bowel disease (9) and multiple cancers (10, 11). Like other chemokine receptors, agonist-mediated activation of CCR5 results in coupling to the cAMP-inhibitory (Gi) class and phospholipase activating (Gq) class of heterotrimeric G proteins (12). Stimulation of PI3K (13), MAPK (14) and JAK/STAT (15) signaling pathways has also been reported. Some ligands also promote phosphorylation of the receptor through G protein-coupled receptor kinases (GRKs) with subsequent engagement of β-arrestins, which regulate signaling through receptor desensitization and internalization (13, 16–18).

Experiments with N-terminally modified CCL5 variants demonstrated that the exact pharmacological responses of CCR5, including its post-activation intracellular trafficking behavior, can be markedly different depending on the nature of the ligand. For example, while WT CCL5 allows for efficient receptor recycling back to the cell surface following initial internalization (16), the superagonist CCL5 variant AOP-CCL5 was shown to induce more rapid and efficient receptor internalization and retention in endosomes: this unique property makes AOP-CCL5 a particularly potent inhibitor of HIV entry into cells (19). These findings motivated systematic efforts to identify additional CCL5 variants with a wide range of pharmacological properties including non-signaling antagonists, superagonists with powerful receptor sequestering properties, as well as ligands with G protein or β-arrestin subtype bias and the ability to induce distinct trafficking patterns (20–23).

Despite published insights into the molecular basis of CCR5 ligand-dependent signaling and trafficking (16, 20, 22, 23), the molecular picture is far from complete. Because of the therapeutic potential for “functional antagonism” (19, 24) that hinges on CCR5 internalization and sequestration, we were particularly interested in understanding the trafficking profiles induced by different ligands and the mechanisms driving this behavior. As receptor trafficking is determined by distinct ligand-dependent interactions with intracellular proteins, we turned to ascorbate peroxidase- (APEX2-) catalyzed proximity biotinylation coupled with mass spectrometry to reveal these interactions and their ligand dependencies. This method enables the identification of directly and indirectly interacting proteins with a high degree of spatial (<20 nm) and temporal resolution when the peroxidase enzyme is fused to the C-terminus of the receptor of interest, as demonstrated for other GPCRs (25–27). Based on patterns of interactors, it can also be used to obtain information on the time-dependent location of the receptor in the cell, which was the primary goal of the present study in order to better understand ligand-dependent sequestration.

We conducted the APEX studies with three chemokine variants that differ only in the first few N-terminal amino acid residues yet induce distinct signaling and trafficking patterns of CCR5 (22): WT CCL5 (CCL5), the super-agonist [6P4]CCL5 (6P4), and [5P14]CCL5 (5P14), a partial and possibly Gi-biased agonist (22). Results from the proximity labeling studies were then combined with a variety of cell-based assays (e.g. immunoprecipitation, flow cytometry, bioluminescence resonance energy transfer (BRET) and fluorescence microscopy) to obtain insight into the ability of these ligands to regulate the kinetics and localization of CCR5 into different intracellular compartments. In addition to its role in migration, CCR5 also scavenges chemokines from the extracellular environment (28). The molecular mechanisms that drive scavenging vs G protein coupling are poorly understood but hypothesized to be critical for maintaining appropriate extracellular chemokine levels, thereby preventing receptor downregulation and regulating the resolution of the inflammatory processes (29–31). As scavenging critically depends on receptor trafficking behavior (30, 31), we were able to correlate the differential ability of CCR5 to scavenge WT, 5P14 and 6P4 with the receptor trafficking profiles induced by these ligands. Overall, the results demonstrate how pharmacologically distinct ligands can markedly alter the endocytic fate of CCR5 through internalization, recycling, degradation and subcellular compartment trapping. The identification of such mechanisms may prove useful for therapies based on exploiting receptor sequestration.

## Results

### Time-resolved ligand-dependent trafficking of CCR5 revealed by APEX2 proximity labeling proteomics

To investigate the ligand-specific trafficking of CCR5, we utilized APEX2 proximity labeling proteomics (25, 26), an unbiased approach that can reveal protein networks in proximity to CCR5 under different stimulation conditions and at different time points. These experiments were conducted in HEK293T cells stably expressing CCR5 C-terminally tagged with APEX2 peroxidase (CCR5-APEX). We also included spatial references of plasma membrane, endosomes and cytosol (see Methods) used in previous APEX studies (25) for distinguishing bystander proteins at those locations from potential interactors of CCR5. Because C-terminal modifications can alter signaling and trafficking of GPCRs (25), we first confirmed that WT CCR5 and CCR5-APEX are similar in their ability to trigger G protein activation and β-arrestin2 recruitment following chemokine stimulation (**Figure S1A-C**); this was done using Bioluminescence Resonance Energy Transfer (BRET)- based G protein dissociation and β-arrestin2 association assays, respectively. Importantly, CCR5-APEX internalized at a level comparable to WT CCR5 after chemokine treatment (**Figure S1D**). We also demonstrated that the three different chemokines under study (CCL5, 6P4, 5P14) induce different signaling and trafficking phenotypes of both WT CCR5 and CCR5-APEX (**Figure S1**). Consistent with previous reports that 6P4 is a super-agonist, it induced greater G_i_ protein activation, β-arrestin2 recruitment and receptor internalization than WT CCL5 at the same concentration (100 nM). By contrast, 5P14 treatment resulted in G_i_ protein dissociation comparable to WT CCL5, but it induced the recruitment of only ∼20% of the β-arrestin2 recruited to the receptor by WT CCL5 (**Figure S1C**). Overall, the results suggest that despite the C-terminal tag, CCR5-APEX faithfully represents WT CCR5, including its differential responses to the three ligands, and is therefore suitable for the proximity labeling experiments.

The ligand treatment scheme and proximity labeling strategy coupled with tandem mass tag mass spectrometry (32) are illustrated in **Figure S2**. 21,550 individual peptides, representing a total of 3,187 proteins, were identified from the entire dataset (see Methods and **Figure S3**). Of these proteins, 428 showed significant variation across all ligand-time conditions **(Figure 1A and Table S1)**, as evaluated by two-way moderated ANOVA with Benjamini-Hochberg (BH) adjustment for multiple comparisons (33) (see Methods). Among the significantly varied proteins, a strong over-representation of Gene Ontology (GO) (34, 35) terms associated with cellular trafficking, signaling, and cytoskeleton and cell-cell junction regulation was observed (**Figure 1B**).

**Figure 1.**
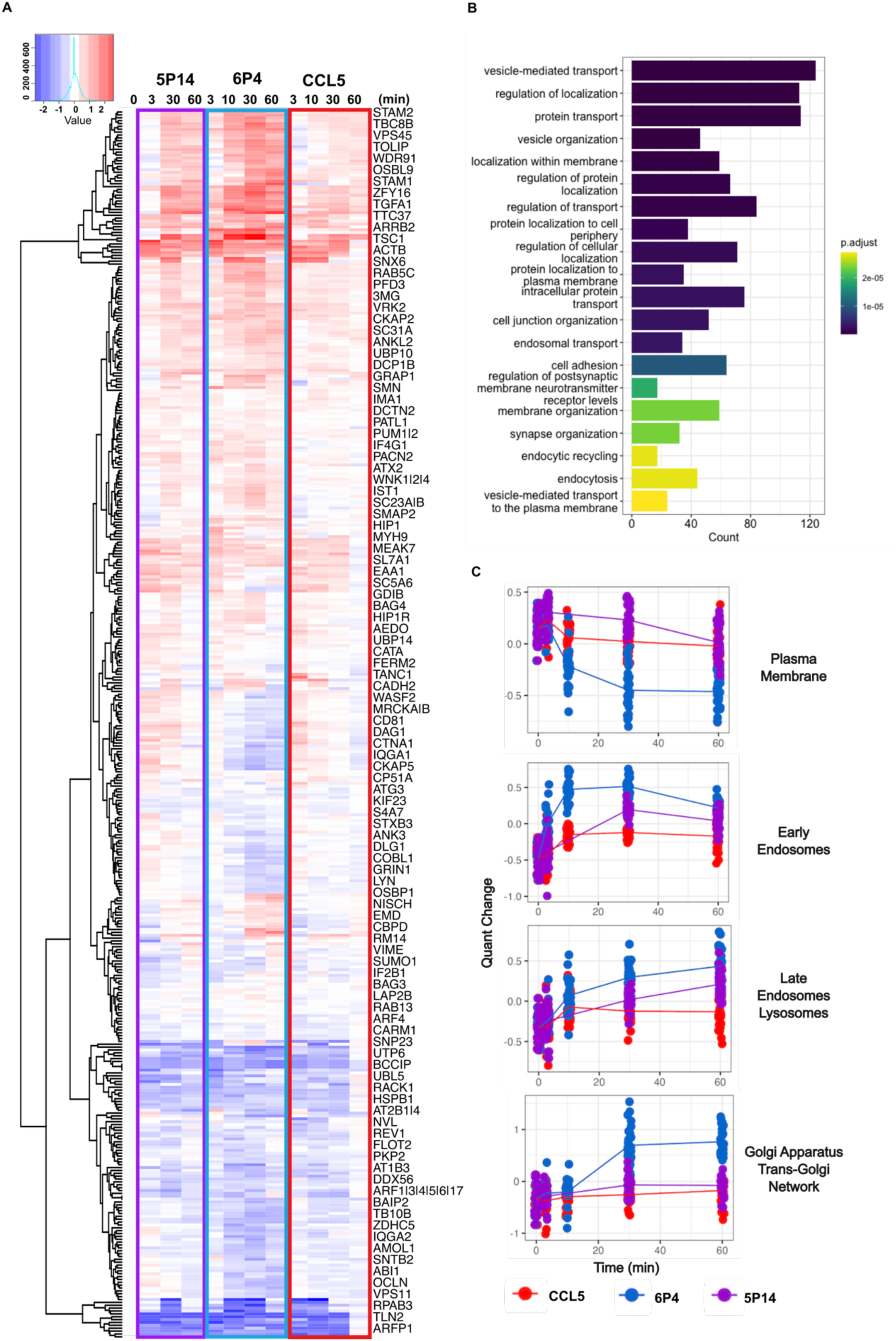
Time-resolved ligand-dependent trafficking of CCR5 revealed by APEX proximity labeling proteomics. (A) Heatmap of time-dependent CCR5-APEX labeling of proteins that significantly varied across all ligand conditions. Protein quants from all ligand conditions (different chemokines at 3, 10, 30 or 60 minutes after stimulation) were normalized to buffer control without stimulation (0 minute). An increase in quants is indicated by varying shades of red corresponding to the level of increase. Decreases in quants are colored in blue with varying shades indicating different levels of decrease. (B) Top 20 overrepresented Gene Ontology (GO) Biological Process (BP) terms among 428 proteins labeled by CCR5-APEX that significantly varied across all ligand conditions. (C) Centers of representative clusters of proteins labeled by CCR5-APEX: response profiles to CCL5, 6P4, 5P14 are shown in red, blue, and purple, respectively. Dots represent aggregated peptide centers from all proteins in the cluster that share the same overall trend in labeling change over time and the line connects the quant distribution center of all members at each timepoint. GO overrepresentation analysis was performed on each cluster and the most significantly overrepresented Cellular Component (CC) term associated with the corresponding cluster is shown on the right.

To further investigate the ligand-dependent trafficking patterns of CCR5, we examined the collective spatiotemporal behavior of the labeled proteins. Specifically, proteins were hierarchically clustered based on their time-dependent labeling patterns, after which over-representation of Gene Ontology Cellular Compartment terms was calculated separately for each cluster (see Methods). The resulting compartment-specific quant profiles demonstrate that agonist-stimulated CCR5 departs from the plasma membrane and traffics to early endosomes, and with slower kinetics, associates with endo-lysosomes, thus following the canonical endosomal trafficking pathway. However, the three ligands induced dramatically different levels and rates of CCR5 internalization, as revealed by the time-dependent variations of proteins in the plasma membrane cluster (**Figure 1C**). For example, 6P4 triggered a significantly greater and more rapid decrease in PM protein labeling compared to WT CCL5, corresponding to its faster and higher level of receptor internalization, whereas 5P14 induced a lower level of change that was more gradual, also consistent with its receptor internalization kinetic profile (**Figure 1C)**. Conversely, most early endosomal proteins exhibited a corresponding increase in labeling over time, indicating accumulation of the receptor in the early endosomal compartments after ligand stimulation, with similar ligand-dependent differences as for the plasma membrane proteins (**Figure 1C**). In addition to these two subcellular locations, we also observed marked ligand-dependent labeling pattern differences for protein clusters associated with the cytoskeleton, late endosomes, lysosomes, endoplasmic reticulum, Golgi, trans-Golgi network, nucleus and exocytic vesicles (**Figure 1C and Table S2**). It is also important to note that our analysis generated not a one-to-one but rather a many-to-many correspondence between clusters and subcellular locations: some locations were overrepresented in more than one response profile cluster and, conversely, most clusters showed over-representation of proteins from more than one subcellular location. For example, plasma membrane proteins were overrepresented in at least 5 clusters, all exhibiting similar overall trends but with different ligand responses (**Table S2**). Interestingly, among them, clusters with overrepresentation of cytoskeletal proteins showed much less variation among ligand conditions compared to the others (**Table S2**). This demonstrates that our APEX proximity data has sufficient spatial resolution to distinguish distinct regions, substructures, or microdomains of a single subcellular location. Overall, this provided a systematic view of the ligand-dependent subcellular trafficking patterns of CCR5 and guided the follow-up studies described in the following sections.

### β-arrestin-dependent clathrin-mediated endocytosis drives ligand-dependent internalization

Ligand-dependent patterns of CCR5 internalization observed by proximity labeling were next validated by a bystander BRET assay measuring proximity between CCR5 C-terminally tagged with *Renilla* luciferase (CCR5-RlucII) and a plasma membrane marker (the polybasic prenylated CAAX box of KRas N-terminally tagged with *Renilla* green fluorescent protein, rGFP-CAAX), or early endosome marker (the rGFP-tagged FYVE domain of endofin, rGFP-FYVE) (36). Chemokine agonists triggered CCR5 departure from the plasma membrane, as demonstrated by a decrease in the BRET ratio between CCR5-RlucII and rGFP-CAAX (**Figure 2A**). Accumulation of the receptor in the early endosome was indicated by an increase in the BRET ratio between CCR5-RlucII and rGFP-FYVE (**Figure 2A**). The different levels and rates of BRET changes associated with each ligand condition matched the ligand-dependent changes in plasma membrane or early endosome protein labeling observed in the APEX data (**Figure 1B**, **Figure 2A and Figure S4A**).

**Figure 2.**
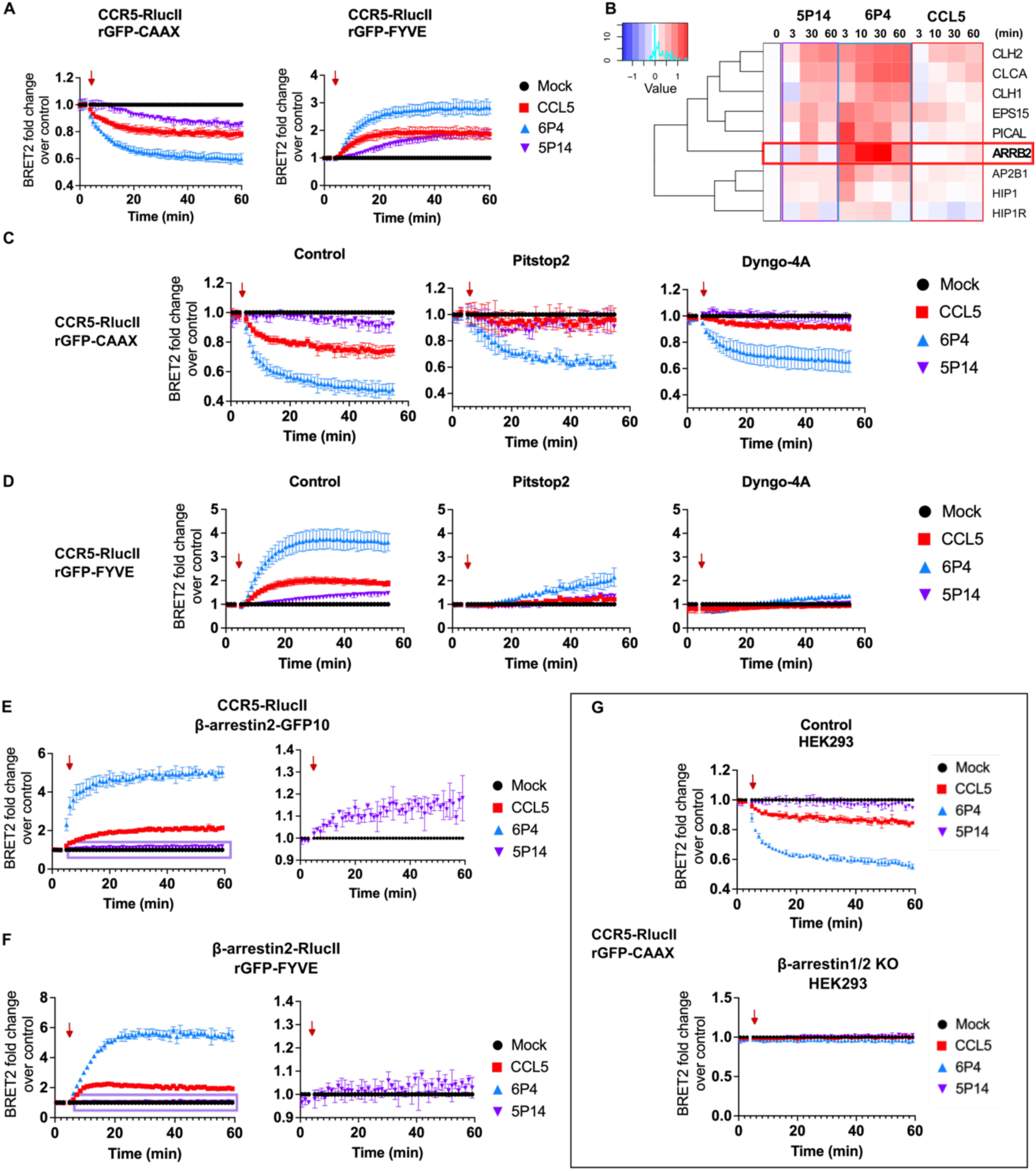
CCR5 internalization by all three ligands is dependent on β-arrestin, proceeds through clathrin-mediated endocytosis, and involves receptor translocation to early endosomes. (A) Ligand-dependent temporal profiles of bystander BRET between CCR5-RlucII and rGFP-CAAX (a PM marker) or rGFP-FYVE (an EE marker) in HEK293T cells support CCR5 ligand-dependent internalization patterns revealed by APEX proteomics. BRET signals from ligand-treated samples (100 nM CCL5, 6P4, or 5P14) are subtracted from buffer-treated controls and are represented as mean ± SE from three independent experiments. Control-subtracted BRET traces from CCL5-, 6P4- and 5P14-treated samples are colored in red, blue and purple respectively; buffer controls are colored in black. Addition of ligands is indicated with a red arrow on the plots. (B) Heatmap of time-dependent CCR5-APEX labeling of proteins associated with clathrin-coated pits and β-arrestin2 (ARRB2, indicated by a red box). (C and D). Effect of clathrin-mediated endocytosis (CME) inhibitors Pitstop^®^2 and Dyngo-4a on ligand-induced internalization of CCR5. (E) BRET between CCR5-RlucII and β-arrestin2-GFP10 in HEK293T cells showing β-arrestin2 recruitment directly to CCR5 after ligand treatment (100 nM CCL5, 6P4, or 5P14). A zoom of the BRET signal trace from the 5P14 treated sample (purple box) is shown on the right. (F) Bystander BRET between RlucII-β-arrestin2 and rGFP-FYVE in CCR5 expressing HEK293T cells showing β-arrestin2 recruitment to early endosomes after ligand stimulation of CCR5 (100 nM CCL5, 6P4, or 5P14). A zoom of the BRET signal trace from the 5P14 treated samples (purple box) is shown on the right. (G) Role of β-arrestin1/2 in ligand-induced internalization of CCR5 revealed by bystander BRET between CCR5-RlucII and rGFP-CAAX in control HEK293 cells or β-arrestin1/2 knockout (KO) HEK293 cells.

After agonist stimulation, CCR5 is known to interact with β-arrestin1/2 and internalize via clathrin-mediated endocytosis (CME) (37). We speculated that β-arrestin1/2 and CME regulators might also be responsible for the ligand-dependent differences in CCR5 internalization. Consistent with our hypothesis, the labeling level and kinetics of β-arrestin2 (ARRB2) as well as proteins known to be involved in CME, such as clathrin light chain and heavy chain proteins, components of AP-2 complex, and Huntingtin-interacting proteins, matched the ligand-dependent CCR5 internalization level and kinetics (**Figure 2B**). Compared to other ligands, 6P4 induced the strongest labeling of β-arrestin2 and known CME regulators, including the phosphatidylinositol-binding clathrin assembly protein PICAL (38) and EGFR substrate EPS15 (39) as early as 3 minutes after stimulation (**Figure 2B**). To further test whether CME is the major pathway for ligand-induced internalization, we used two different pharmacological inhibitors of CME: pitstop2^®^ and Dyngo-4A. The former inhibits interactions between CME regulators and clathrin (40) and the latter inhibits dynamin-dependent membrane fusion (41). Both inhibitors significantly suppressed CCR5 internalization by all three ligands as shown by the CCR5-RLucII/rGFP-CAAX bystander BRET results (**Figure 2C and Figure S4B)**; they also almost completely abrogated receptor entry into early endosomes (**Figure 2D and Figure S4C**). Interestingly, CCR5 internalization by CCL5 and 5P14 was more severely inhibited, whereas its internalization by 6P4 was partially preserved. This may reflect an incomplete blockade of CME by pharmacological inhibitors.

Since β-arrestins have been shown to play important roles in CCR5 internalization (18) as well as more generally in internalization of GPCRs (42, 43), we further investigated the interaction of CCR5 with β-arrestin1/2 following the different ligand treatments. As mentioned above, 6P4 induced the fastest, strongest, and most persistent labeling of β-arrestin2. CCL5 on the other hand, induced much lower and more gradual labeling compared to 6P4, but the labeling was still persistent, with appreciable signal even at 60 minutes. However, with 5P14 treatment, β-arrestin2 labeling was not only slower, but more transient as well; it exhibited a peak at 30 minutes but dropped significantly by 60 minutes (ARRB2 in **Figure 2B**). Again, the ligand differences in proximity labeling patterns were corroborated by BRET-based effector membrane translocation assays using two different donor-acceptor pairs: (i) CCR5-RlucII and C-terminally GFP10-tagged β-arrestin2 (β-arrestin2-GFP10), which monitors the direct recruitment of β-arrestin2 to the receptor (44) and (ii) C-terminally RlucII-tagged β-arrestin2 (β-arrestin2-RlucII) and rGFP-FYVE, which monitors the translocation of β-arrestin2 to the early endosome (45). Recruitment of β-arrestin2 to the receptor or to the early endosome compartment was demonstrated by increases in BRET ratios of these donor-acceptor pairs. Distinct levels and kinetics for each ligand were observed, consistent with the proximity labeling data (**Figure 2E and Figure S4D**). Importantly, we demonstrated robust recruitment of β-arrestin2 to the early endosomal compartment with 6P4 and CCL5 treatment; however, with 5P14, such recruitment was virtually undetectable (**Figure 2F**). Since β-arrestin2 recruitment and CCR5 entry to the early endosome both occurred without delay and with similar kinetics, it is likely that β-arrestin2 accompanied the receptor from the plasma membrane to the early endosome after CCL5 and 6P4 treatment as opposed to direct recruitment of β-arrestin2 to the receptor at the early endosome. This is consistent with the ability of CCL5 and 6P4 to induce more persistent β-arrestin2 interactions with CCR5, whereas 5P14 only induces transient interactions, as reported previously (21). Because in HEK293T cells, β-arrestin1 is much less abundant than β-arrestin2, our mass spectrometry experiment did not identify any endogenous β-arrestin1 peptides in CCR5-APEX-labeled samples; however, β-arrestin1 was similar to β-arrestin2 in the corresponding BRET-based assays (**Figure S5**).

To investigate the importance of β-arrestin1/2 to ligand-induced internalization of CCR5, we next assessed CCR5 departure from the plasma membrane (via CCR5-RluII/rGFP-CAAX bystander BRET) in a β-arrestin1/2 CRISPR knock out (KO) HEK293 cell line. Strikingly, knocking out β-arrestin1/2 almost completely eliminated ligand-dependent CCR5 internalization, even with 6P4 treatment (**Figure 2G and Figure S4E**). Altogether, these data indicate that CCR5 internalization by all three ligands is β-arrestin-dependent and the internalization is primarily clathrin-mediated.

### CCR5 phosphorylation by different GRK subtypes affects ligand-dependent internalization through β-arrestin1/2

It is well established that phosphorylation of GPCRs by GPCR-kinases (GRKs) is a prerequisite for β-arrestin recruitment and β-arrestin-dependent internalization (46). We therefore investigated whether CCR5 phosphorylation by different GRK subtypes impacts ligand-induced internalization and explains at least in part the ligand-dependent differences observed in the β-arrestin2 interactions with CCR5. To this end, we conducted CCR5-RlucII and rGFP-CAAX BRET experiments in three HEK293 cell lines in which different combinations of GRKs were knocked out (ΔGRK2/3, ΔGRK5/6, and ΔGRK2/3/5/6). In the ΔGRK2/3 HEK293, CCL5-mediated internalization of CCR5 was almost completely inhibited, whereas knock out of GRK5/6 (ΔGRK5/6) had no effect (**Figure 3A and Figure S6A**) indicating the dominance of the Gβγ-dependent GPCR kinase(s). The signal of 5P14-induced internalization was small in parental HEK293 cells so the differences among GRK knockout cell lines were harder to interpret (**Figure 3A and Figure S6A**). Surprisingly, neither GRK2/3 or GRK5/6 knock out had a large impact on internalization induced by 6P4 (**Figure 3A and Figure S6A**). Contrary to the diverse effects of double-GRK knockouts, the elimination of all four GRKs (ΔGRK2/3/5/6 HEK293) completely abrogated CCR5 internalization by all three ligands (**Figure 3A and Figure S6A**), consistent with its dependence on receptor interaction with β-arrestin1/2. Furthermore, the complete elimination of 6P4-induced receptor internalization in the ΔGRK2/3/5/6 cells but the lack of an effect in the ΔGRK2/3 and ΔGRK5/6 cells indicates that with 6P4-stimulated receptor, there is compensation by the remaining GRKs in these latter two knockout cell lines.

**Figure 3.**
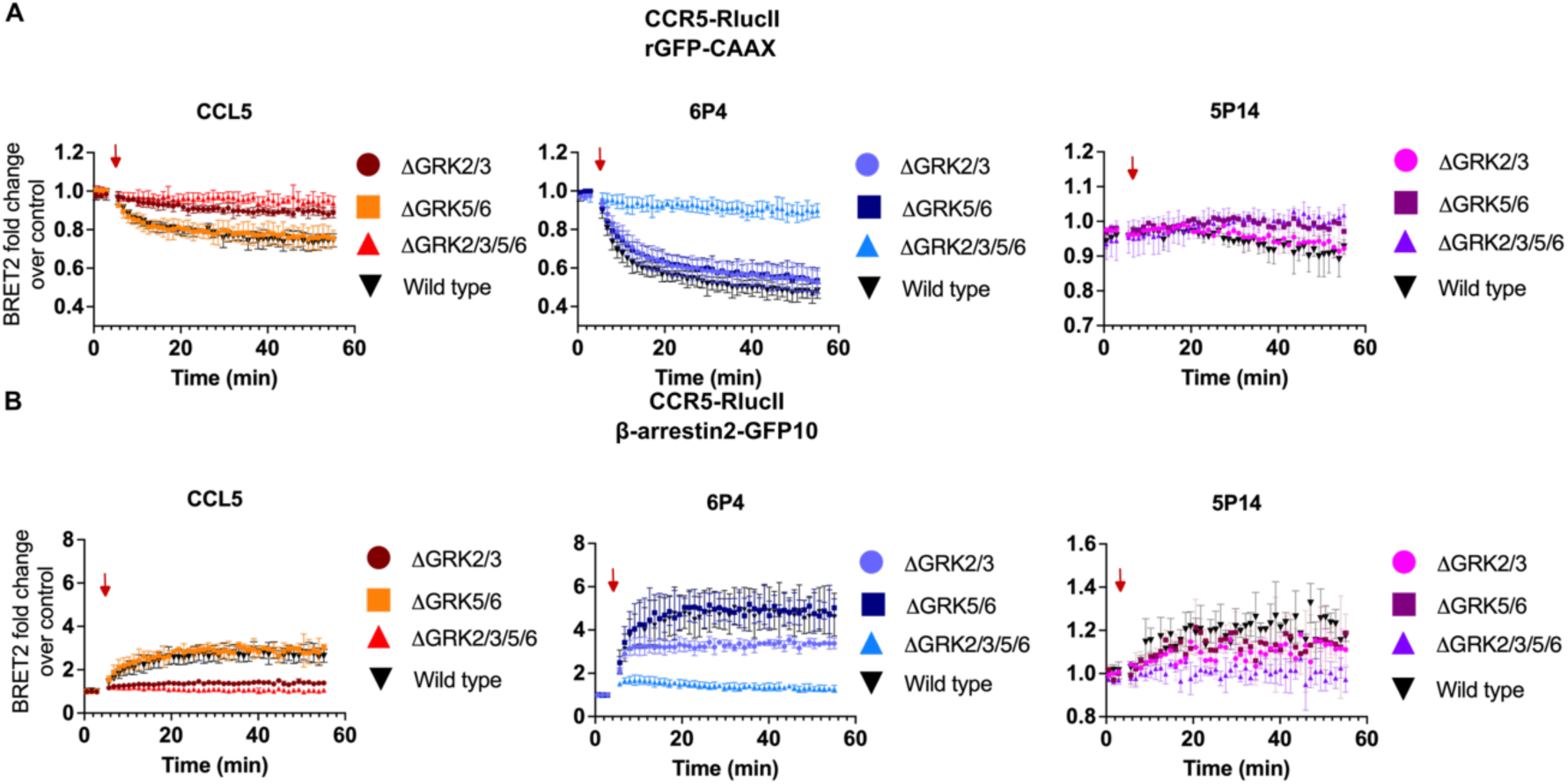
CCR5 phosphorylation by different GRK subtypes affect ligand-dependent internalization through β-arrestins recruitment. (A and B) BRET between CCR5-RlucII and rGFP-CAAX (PM marker, A) or β-arrestin2-GFP10 (B) in parental HEK293A cells, ΔGRK2/3, ΔGRK5/6, or ΔGRK2/3/5/6 knockout HEK293A cells. BRET signals from ligand-treated groups are subtracted from buffer-treated controls and are represented as mean ± SE from three independent experiments. Control-subtracted BRET traces of different cellular backgrounds from the same ligand treatment condition (100 nM CCL5, 6P4, or 5P14) are plotted together to better demonstrate the differences in CCR5 PM departure (A) or β-arrestin2 recruitment (B) in the GRK knockout lines.

To examine the dependence of the CCR5 interaction with β-arrestin2 on the different GRKs, we directly measured β-arrestin2-GFP10 association with CCR5-RlucII by BRET in the three GRK subtype knockout cell lines. With CCL5 stimulation, β-arrestin2 recruitment to CCR5 was severely impacted in ΔGRK2/3 but not in the ΔGRK5/6 cells (**Figure 3B and Figure S6B**). With 5P14 treatment, similar to internalization, the β-arrestin2 recruitment to CCR5 was at a much lower level even in the parental HEK293 cells, making comparisons in the GRK knockout inconclusive (**Figure 3B and Figure S6B**). With 6P4 stimulation, β-arrestin2 recruitment was inhibited in the ΔGRK2/3 but not in the ΔGRK5/6 cells, although the defect in ΔGRK2/3 cells was not as severe as for the WT CCL5-stimulated receptor and it did not negatively impact receptor internalization (**Figure 3B and Figure S6B**); this suggests again that GRK2/3 is the dominant kinase although GRK5/6 can compensate in the absence of the G protein-dependent (GRK2/3) kinases. As expected, agonist-induced β-arrestin2 recruitment to CCR5 was completely inhibited in the ΔGRK2/3/5/6 cells regardless of the agonist variant used (**Figure 3B and Figure S6B**). Overall, the patterns of ligand-dependent β-arrestin2 recruitment to CCR5 tracked well with its ligand-dependent internalization (**Figure 3B**). Together these data suggest that the different ligands induce different levels and/or patterns of CCR5 phosphorylation by GRKs, which in turn affect its ability to recruit β-arrestin1/2 and internalize.

### The CCL5 ligands differentially affect receptor turnover at the cell surface but do not impair recycling pathways

Some of the ligand- and time-dependent proximity biotinylation profile clusters in our data show over-representation of proteins from subcellular locations other than PM or early endosomes (**Figure 1B and Table S1**). This indicates that the ligands differentially regulate not only CCR5 internalization but also other aspects of CCR5 trafficking inside the cell. Consistent with this, previous research demonstrated prolonged intracellular receptor retention following treatment with 6P4 and 5P14 but not CCL5 (22), indicating potential effects on recycling. To further probe the mechanisms, we first assessed the constitutive and ligand-induced internalization of CCR5 via pre-label vs post-label flow cytometry (see Methods). In the pre-label assay, cell surface receptors are labeled with a primary antibody <u>before</u> allowing them to internalize; then, after a defined period of incubation at 37°C, receptors remaining on the cell surface are quantified with a secondary antibody. Neither the receptors that internalized nor those that reappear on the cell surface via recycling are detected by the secondary antibody. By contrast, in the post-label assay, both the primary and the secondary antibody are applied <u>after</u> allowing receptors to internalize for a given period of time at 37°C. This protocol ensures the detection of all cell surface receptors, including those that remain on the surface after internalization and those that emerge on the surface as a result of recycling. Thus, the difference in the surface level of receptor between the pre-label and the post-label assay indicates the extent of receptor recycling.

Based on results of the pre-label experiment, 1 hr treatment with CCL5 and 5P14 caused internalization of approximately 65% and 60% of CCR5 (**Figure 4A**). Using 6P4, almost 90% of the receptor was internalized **(Figure 4A).** Note that CCR5 also constitutively internalizes as indicated by the similar level of internalization of the receptor under buffer only conditions (indicated as “mock” in **Figure 4A**) and induced by WT CCL5. As indicated by the differences in the fluorescent staining between the pre-labeled and post-labeled cells, the extent of CCR5 recycling to the surface was approximately 25% for CCL5, but only 10% in the 6P4-treated samples and 6% in the 5P14-treated samples after 30 min (**Figure 4A**). As additional evidence, we evaluated CCR5-RlucII re-association with the plasma membrane (via the CCR5-RlucII/rGFP-CAAX BRET assay). In these experiments, cells were treated with the CCR5 antagonist Maraviroc after 30 minutes of agonist-induced internalization to competitively inhibit receptor interaction with chemokine and thus prevent further chemokine-induced internalization (31). Interestingly, 6P4-treated CCR5 returned to the surface at a higher level compared to WT CCL5 (**Figure 4B)**. However, when normalized to the total amount of internalized CCR5, the percentage of the recycling receptors was significantly lower following 6P4 treatment **(Figure 4C)**. Consistent with the results from the pre- and post-label experiments, 5P14-treated receptor showed much slower internalization kinetics and reduced recycling to the cell surface compared to the other two ligands (**Figure 4B-C**). These results are consistent with previous research that showed more retention of CCR5 inside the cells when stimulated by 6P4 and 5P14 (22).

**Figure 4.**
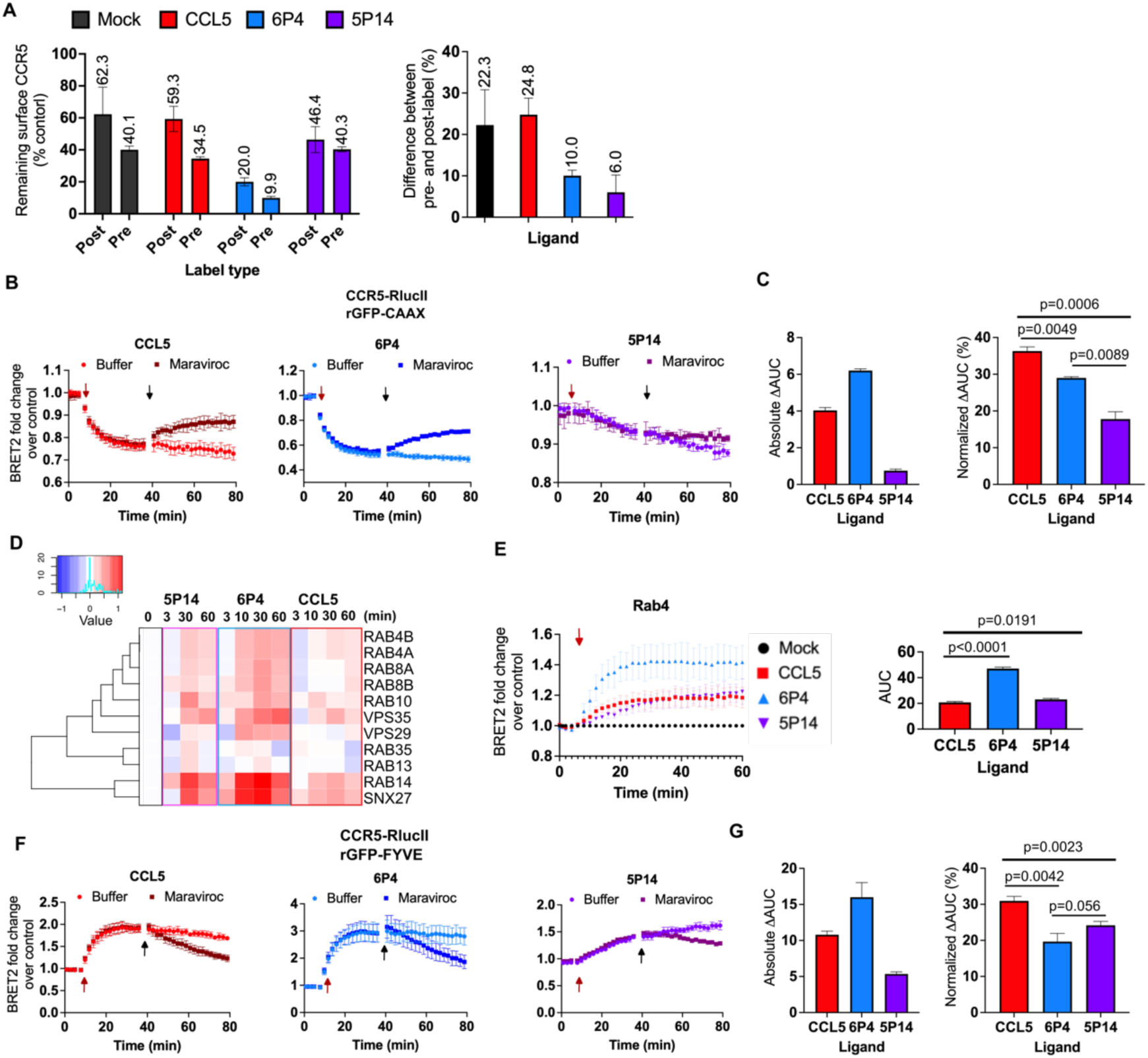
The CCL5 ligands differentially affect receptor turnover at the cell surface but do not impair recycling pathways. (A) Internalization of HEK293 cells stably expressing WT CCR5 after ligand stimulation (100 nM CCL5, 6P4, or 5P14) were assessed by pre-label or post-label flow cytometry (see Methods). On the right, the remaining CCR5 surface level difference between pre-label and post-label assays is presented for each ligand condition. Data are presented as a fraction of surface receptor remaining compared to non-internalized control at 4 °C. (B) Bystander BRET between CCR5-RlucII and rGFP-CAAX in cells stimulated with the indicated agonists (red arrow) for 30 min, followed by the addition of 5uM Maraviroc^®^ (black arrow). BRET signals from ligand-treated samples are subtracted from buffer-treated controls and are represented as mean ± SE from three independent experiments. Control-subtracted BRET traces of either inhibitor- or buffer-treated samples from the same ligand treatment condition are plotted together to demonstrate the differences in the BRET signal after antagonist addition. (C) Area under curve (AUC) was calculated for BRET traces in (B) with the baseline set to 1. The differences between the AUC of the buffer and the Maraviroc^®^ treated samples is plotted for each ligand (left) and the normalization of the difference against the AUC of the buffer control for the corresponding ligand is plotted together for comparison (right). All data are expressed as the mean ± SE of n = 3 independent experiments. P values were calculated with unpaired t test with Welch’s correction. (D) Heatmap of time-dependent CCR5-APEX labeling of proteins associated with recycling pathways. (E) Bystander BRET between CCR5-RlucII and rGFP-Rab4 (a fast-recycling endosome marker) in HEK293T cells treated with 100 nM CCL5, 6P4, or 5P14. AUC quantification is presented on the right. All data are expressed as the mean ± SE of n = 3 independent experiments. P values were calculated with the unpaired t test with Welch’s correction. (F) Bystander BRET between CCR5-RlucII and rGFP-FYVE in cells stimulated with the indicated agonists (red arrow) for 30 min, following the addition of 5 uM CCR5 inhibitor Maraviroc^®^ (black arrow). BRET signals from ligand-treated samples are subtracted from buffer-treated controls and are represented as mean ± SE from three independent experiments. Control-subtracted BRET traces of either inhibitor- or buffer-treated samples from the same ligand treatment condition are plotted together to better demonstrate the differences in BRET signal after antagonist addition. (G) AUC was calculated for BRET traces in (F) with the baseline set to 1. The differences between the AUC of the buffer and the Maraviroc^®^ treated samples are plotted for each ligand (left) and the normalization of the difference against the AUC of the buffer control for the corresponding ligand is plotted together for comparison (right). All data are expressed as the mean ± SE of n = 3 independent experiments. P values were calculated with the unpaired t test with Welch’s correction.

Since 6P4 and 5P14 cause CCR5 to be retained inside the cells, we wondered whether they negatively impact receptor association with components of the recycling pathway, compared to CCL5. However, according to the APEX proximity labeling data, 6P4 and 5P14 not only induced labeling of those proteins, but they did so at a comparable or higher level than CCL5 (**Figure 4D**). This was especially the case for Rab4, Rab14, and SNX27, which are involved in sorting and recycling of early endosomes (**Figure 4D**). A BRET-based assessment of CCR5-RlucII proximity to C-terminally rGFP-tagged Rab4 (36) similarly showed higher recruitment of CCR5 to Rab4-positive recycling endosomes when treated by 6P4 compared to 5P14 and CCL5, which were similar to each other (**Figure 4E**). This suggests that 6P4 and 5P14 do not impair the trafficking of CCR5 into receptor recycling pathways; it is also unlikely that they cause CCR5 to be trapped in the early endosomes. Indeed, BRET assays using CCR5-RlucII and rGFP-FYVE showed receptor exiting from early endosomes after Maraviroc treatment with all three chemokines, albeit to a lesser extent after 6P4 or 5P14 treatment (**Figure 4F-G**). This suggests that other mechanisms, described below, may contribute to the intracellular retention of both 6P4 and 5P14-treated CCR5.

### Compared to CCL5, 6P4 and 5P14 induce greater degradation of CCR5 through the endo-lysosomal degradation pathway

Our CCR5-APEX proximity labeling data revealed three different response profile clusters that feature significant overrepresentation of proteins located at late endosomes and lysosomes and higher labeling following 6P4 and 5P14 treatment compared to CCL5 (**Figure 1B and Table S1**). Notable examples within these clusters are late endosomal protein Rab7, lysosomal protein LTOR1, and components of the Endosomal Sorting Complex Required for Transport (ESCRT) pathway such as STAM1/2, HGS, TOM1, TOLIP and Endofin (ZFY16), all of which exhibited higher labeling after 10 min of 6P4 or 5P14 stimulation compared to CCL5 (**Figure 5A)**. A bystander BRET-based assessment of CCR5-RlucII proximity to C-terminally rGFP-tagged Rab7 suggested that CCR5 does enter Rab7-positive late endosomes following treatment with 6P4 and 5P14, but to a lesser extent when treated with CCL5 (**Figure 5B**). Thus, we hypothesized that 6P4 and 5P14 might induce more CCR5 degradation through the endo-lysosomal degradation pathway. To test this, we performed degradation assays using HEK293 cells expressing Flag-tagged CCR5, and indeed observed significantly more receptor degradation following treatment with 6P4 and 5P14 than CCL5 (**Figure 5C**). In support of our hypothesis, this difference was eliminated when the cells were pre-treated with bafilomycin, which inhibits lysosome function by targeting the V-ATPase and preventing lysosomal acidification (47) (**Figure 5C**). No ligand-mediated difference in receptor degradation was observed using the β-arrestin1/2 KO and ΔxGRK2/3/5/6 cells (**Figure 5D**), presumably due to impaired ligand-induced internalization of CCR5.

**Figure 5.**
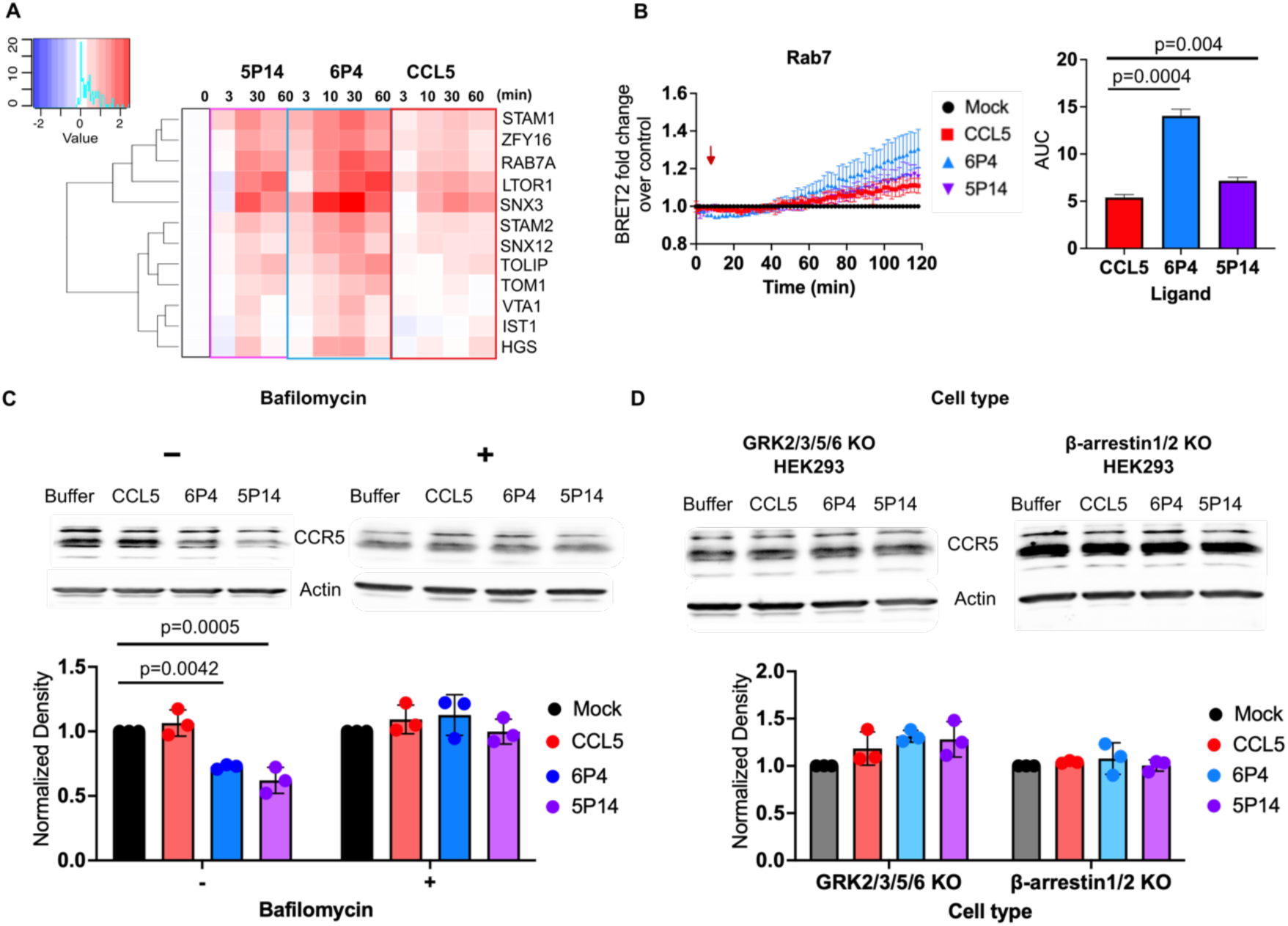
6P4 and 5P14 induce more CCR5 degradation through the lysosomal degradation pathway than CCL5. (A) Heatmap of time-dependent CCR5-APEX labeling of proteins associated with late endosomes, lysosomes and endo-lysosomal degradation pathways. (B) Bystander BRET between CCR5-RlucII and rGFP-Rab7, a late endosome marker, in HEK293T cells stimulated with 100 nM of CCL5, 6P4, or 5P14. AUC quantification is presented on the right. (C) Western blot detection of total Flag-CCR5 in HEK293T cells pretreated or untreated with 1μM bafilomycin (12 hrs), pretreated with 50 μg/mL cycloheximide (15 min), and stimulated or untreated with 100 nM CCL5, 6P4, or 5P14 for 4 hrs at 37°C in the presence of cycloheximide. Endogenous α-actin was used as a loading control. Receptor degradation was quantified, and the data (mean ± SE) shown are expressed as the fraction of receptor remaining compared with untreated control cells as determined from three independent experiments (lower panel). (D) Degradation assays were performed as in (C) but using β-arrestin1/2 KO and GRK2/3/5/6 KO HEK293 cells and no bafilomycin treatment. All data are expressed as the mean ± SE of n = 3 independent experiments. P values were calculated with the unpaired t test with Welch’s correction.

### 6P4 directs CCR5 to the Golgi and Trans-Golgi Network more than CCL5 and 5P14

Several APEX proximity biotinylation profile clusters featured overrepresentation of proteins from the Golgi apparatus and the Trans-Golgi network (TGN), all of which exhibited higher labeling following 6P4 treatment compared to the other ligands (**Figure 1B and Table S1**). One of the clusters showed a marked increase in labeling of Golgi/TGN resident oxysterol-binding proteins and members of the Golgin family (GOGA2-4 and GOGB1) and did so only after 6P4 stimulation for 30-60 minutes **(Figure 1B and Figure 6A)**. This suggests that besides directing CCR5 to late endosomes and lysosomes, 6P4 also directs more receptor to the Golgi apparatus and TGN compared to the other two ligands. To further validate these APEX proximity labeling observations, we assessed SNAP-CCR5 co-localization with a TGN marker (Trans-Golgi network integral membrane protein 2, TGOLN2, or TGN46) by confocal microscopy in HEK293T cells. The microscopy images showed a strong co-localization of CCR5 with the TGN marker only after 6P4 treatment, as indicated by a significantly higher Pearson’s correlation coefficient value compared to other conditions, whereas the internalized CCR5 after 5P14 and CCL5 treatment did not substantially accumulate in the TGN **(Figure 6B-C).** Taken together, it appears that 6P4 sequesters more CCR5 inside the cells for a prolonged period compared to other CCL5 ligands by mechanisms involving degradation and/or by directing CCR5 to the Golgi and TGN, whereas 5P14 retains CCR5 by receptor degradation and/or inducing slower receptor internalization and recycling.

**Figure 6:**
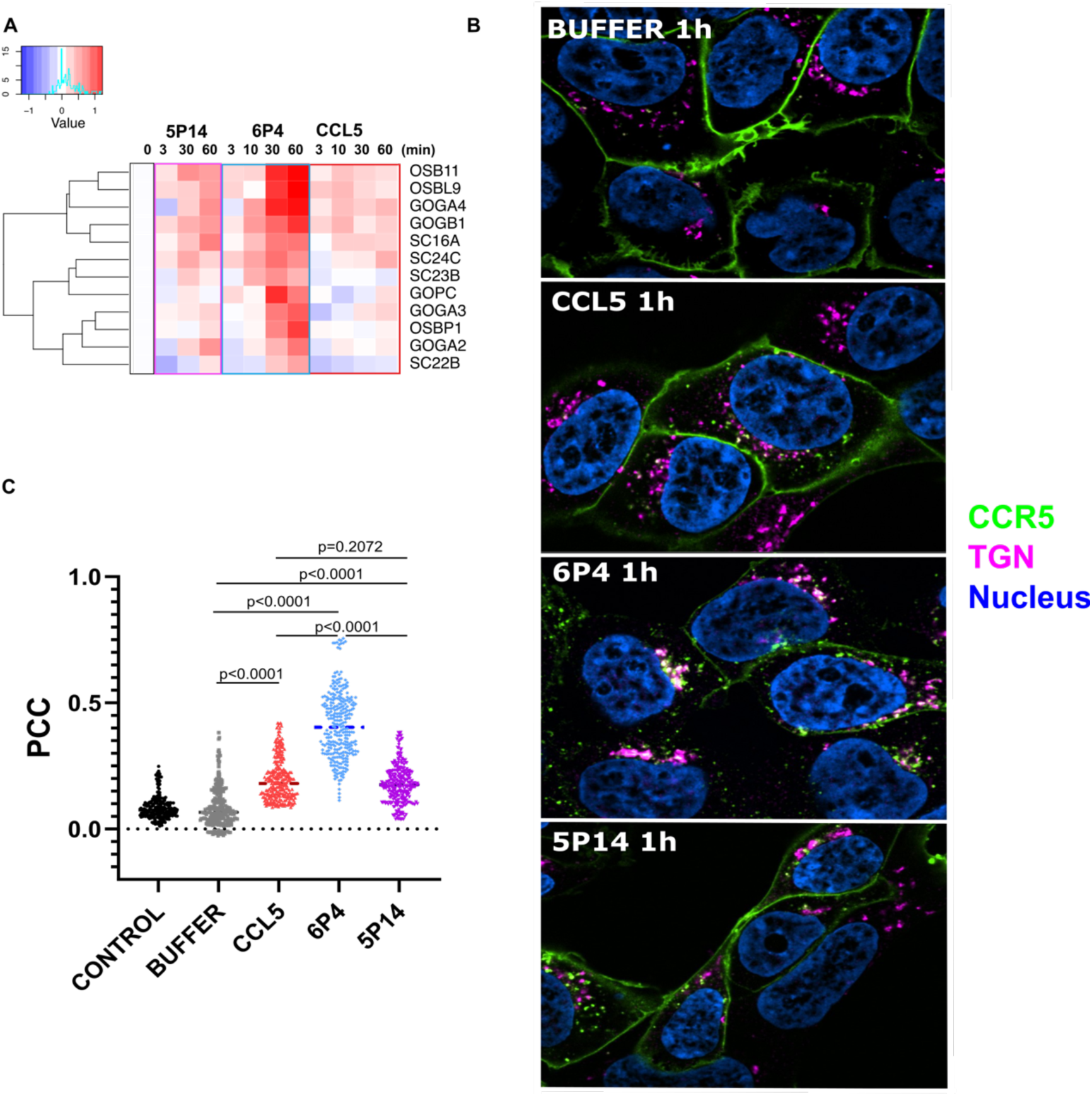
6P4 directs CCR5 to Golgi and TGN more than CCL5 and 5P14. **(A)** Heatmap of time-dependent CCR5-APEX labeling of proteins associated with the Golgi apparatus and Trans-Golgi Network (TGN). (B) Representative microscopy images showing HEK293 cells expressing SNAP-CCR5 labeled in green with TGN markers labeled in magenta; 1 h treatment with either buffer or 100 nM CCL5, 6P4, 5P14. Images are representatives of n = 3 independent experiments, where > 30 cells were imaged for each condition. (C) Pearson’s correlation coefficient (PCC) was calculated for CCR5 and TGN. More than 200 Z-stack images containing > 30 cells per condition were analyzed. Each dot represents the PCC obtained from each Z-stack image. ****P<0.0001 determined by the Mann–Whitney test.

### CCR5 differentially scavenges CCL5 ligands in a β-arrestin-dependent manner

Previous research has shown that CCR5 scavenges CCL3, CCL4 and CCL5, which is hypothesized to regulate chemokine levels and receptor responsiveness, and dampen inflammatory responses induced by chemokines (28, 48). Chemokine scavenging by receptors is largely driven by internalization of the receptor-chemokine complex, intracellular release and degradation of the chemokine in the lysosomes, and recycling of chemokine-free receptors back to the cell surface to pick up more ligand and repeat the process (29, 31). Since the three CCL5 ligands induce distinct trafficking behaviors of CCR5, we hypothesized that the scavenging efficiency of CCR5 would be ligand dependent. To test this, the CCL5 ligands were incubated with the CCR5-expressing HEK293 cells or non-expressing control cells, and the amount of the ligand remaining in the media after 16 h was quantified by ELISA. As shown in **Figure 7**, while CCR5 scavenged approximately 50% of WT CCL5 relative to control, its ability to scavenge the other two engineered ligands was significantly impaired. Since we previously demonstrated that β-arrestin1/2 is required for ligand-induced receptor internalization, we performed the same experiment in β-arrestin1/2 knock out HEK293 cells and found that the scavenging ability of CCR5 for all three ligands was markedly inhibited (**Figure 7**). The inhibition was almost complete for CCL5 and 6P4 in the knockout cells, whereas inhibition of 5P14 scavenging was less severe (**Figure 7**). This correlates with the apparent strength and persistence of the CCR5 interaction with β-arrestin1/2 following the different ligand treatments. Overall, our data indicate that the ability of CCR5 to scavenge different CCL5 ligands varies in accordance with the trafficking patterns induced by these ligands.

**Figure 7:**
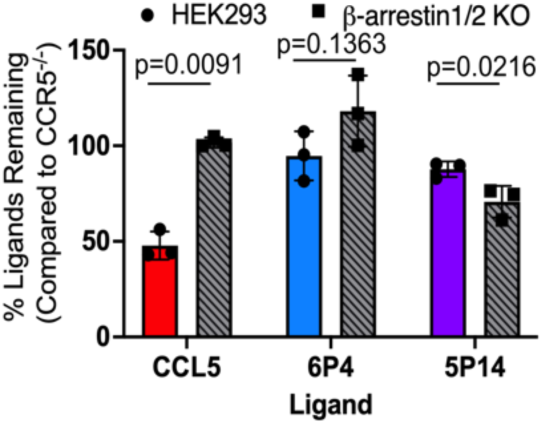
CCR5 differentially scavenges CCL5 variants in a β-arrestin1/2-dependent manner. HEK293 parental and β-arrestin1/2 KO HEK293 cells stably expressing CCR5 and respective non-expressing cells were cultured in media containing 5 nM CCL5 ligands (CCL5, 6P4, or 5P14) for 16 h. Remaining ligands were quantified by ELISA by interpolating from CCL5 standards and presented as a percentage of respective non-CCR5 expressing cells. All data are expressed as the mean ± SE of three independent experiments. P values were calculated using the unpaired t test with Welch’s correction.

### 14-3-3 protein zeta/delta (1433ζ) is a possible contributor to distinct CCR5 intracellular trafficking induced by engineered chemokines

As demonstrated above, the analysis of APEX biotinylated proteins based on statistical significance and collective behaviors (clustering) successfully recapitulated known global trafficking patterns of agonist stimulated CCR5, and added nuances and important distinctions when the receptor is stimulated by WT CCL5 vs its engineered variants. The most pronounced differences were observed with [6P4]CCL5; the trends with [5P14]CCL5 were subtle and the molecular associations that dictate its peculiar effects on CCR5 trafficking remained elusive. We thus asked what specific proteins get recruited to CCR5 stimulated by [5P14]CCL5 but not by the other two chemokines. For this, we selected proteins whose response profiles to CCL5 and [6P4]CCL5 were similar to each other but distinct from [5P14]CCL5, and whose proximity to CCR5 increased (rather than decreased) at both 3 min and 10 min following [5P14]CCL5 stimulation (**Fig. 8A**). Of the nine proteins that met search criteria (**Fig. S8A)**, 14-3-3 protein zeta/delta (1433ζ) attracted our attention as a possible contributor to the observed unique patterns of [5P14]CCL5-stimulated CCR5 intracellular trafficking.

**Figure 8:**
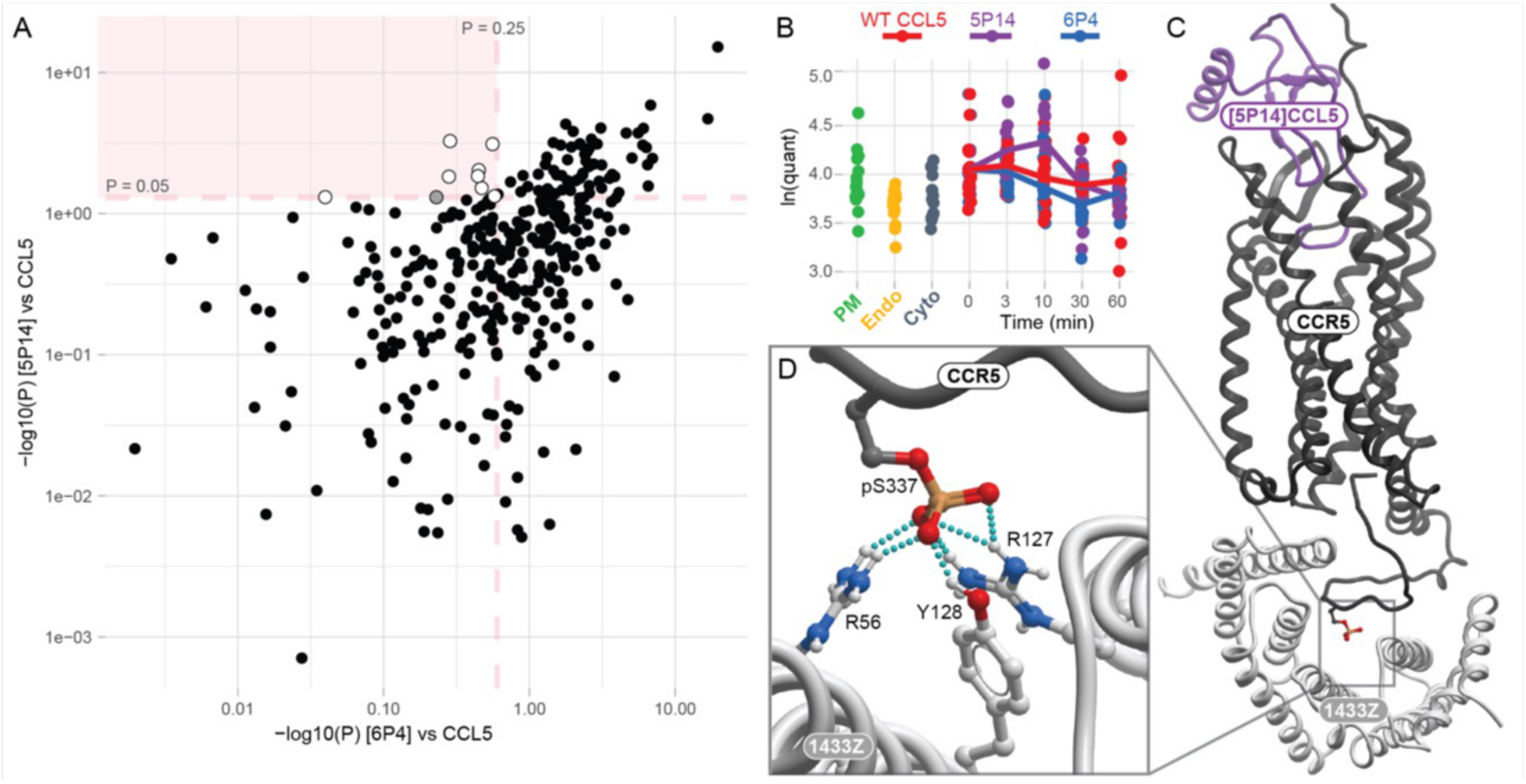
The 1433ζ protein is a potential determinant of unique trafficking of [5P14]CCL5-stimulated CCR5. (**A**) A scatter plot of significance values of [5P14]CCL5 vs [6P4]CCL5 response profile differences relative to WT CCL5, filtered for increase in proximity in both 3 min and 10 min post [5P14]CCL5 addition. The highlighted top left quadrant contains proteins whose [5P14]CCL5 response profile is different from their response to WT CCL5 (P < 0.05), but their response to [6P4]CCL5 is no different from WT CCL5 (P > 0.25); the response profiles of these proteins are shown in **Fig. S8A**. (**B**) The 1433ζ biotinylation temporal profile showed a 0-10 min increase exclusively in response to [5P14]CCL5 and exceeded APEX compartment markers (plasma membrane, endosome, and cytosol). (**C-D**) A 3D model of [5P14]CCL5-bound CCR5 (purple and black, respectively) in complex with 1433ζ (gray). (**C**) The overall view of the complex and (**D**) a close-up of the interaction interface. The C-terminal tail of CCR5 binds in the amphipathic groove of 1433ζ, with CCR5 pSer337 (shown in black sticks) forming six hydrogen bonds with Arg56, Arg127 (both known pSer coordination hubs (67)), and Tyr128 of 1433ζ. Structures were predicted using the 1433ζ homodimer; the second subunit is not shown (see Methods).

In accordance with the search criteria, the proximity of 1433ζ to CCR5-APEX significantly increased when the receptor was stimulated with [5P14]CCL5 but not with WT CCL5 or [6P4]CCL5 (**Fig. 8B**). It then returned to the baseline 30 min after stimulation, also matching the same biotinylation levels as 30 min post CCL5 treatment. In contrast, [6P4]CCL5-triggered biotinylation of 1433ζ decreased for 30 minutes.

1433ζ is one of the seven known isoforms of 14-3-3 proteins, ubiquitous adapters that bind to at least 1% of the mammalian proteome using short linear interacting motifs (SLIMs) sparsely phosphorylated at Ser and Thr residues (49, 50). They have been reported to interact with GPCRs both basally and in an agonist-regulated manner, and to compete with β-arrestins for binding GPCR C-terminal tails (51). We thus hypothesized that the slow and transient 1433ζ recruitment to CCR5 in [5P14]CCL5-stimulated cells is similarly mediated by the C-terminal tail of CCR5, most likely phosphorylated at Ser337 as suggested by the 14-3-3-Pred server (52).

To test this, we constructed AlphaFold3 (53) models of 1433ζ in complex with CCR5 phosphorylated at one of its known C-terminal phosphorylation sites: Ser336, Ser337, Ser342, and Ser349 (54). Despite the modest confidence of these predictions (as evidenced by the pLDDT scores), the C-terminal tail of CCR5 was found to bind to 1433ζ in the canonical geometry in all cases. The complex with CCR5 (pS337) (**Fig. 8C**) featured the best pLDDT scores and conformational consistency across the AF3-predicted model ensemble; we thus hypothesize that Ser337 phosphorylation may mediate the [5P14]CCL5-triggered association between CCR5 and 1433Z is specifically promoted by CCR5 phosphorylation at Ser337 (**Fig. 8D**). As shown in **Fig. 3**, CCR5 internalization in response to WT CCL5 is dependent on GRK2/3 but not on GRK5/6, whereas that induced by [5P14]CCL5 is dependent on GRK5/6 but not on GRK2/3. Therefore, the chemokines can plausibly recruit different GRKs to CCR5 and result in distinct patterns of C-terminal tail phosphorylation. In principle, a sparse or single phosphorylation pattern involving Ser337 may promote CCR5 association with 1433ζ while being insufficient for robust recruitment of β-arrestin. Once bound, 1433ζ can mask the C-terminal tail of CCR5 and keep it away from β-arrestin, explaining why [5P14]CCL5 triggers only minimal β-arrestin association (**Fig. 2E** and **S8B**). Bound 1433ζ may also keep the C-terminus of CCR5 away from the recycling machinery (e.g. SNX27 **Fig. S8C**), thereby possibly explaining why [5P14]CCL5-stimulated CCR5 is preferentially targeted to the late endosomes and lysosomes compared to WT CCL5. Future studies will be focused on testing whether 1433ζ does indeed drive the [5P14]CCL5-specific trafficking of CCR5 compared to the other two ligands.

## Discussion

A major opportunity for developing effective therapeutics against CCR5 is the ability of different ligands to regulate receptor levels on the cell surface. Most efforts to therapeutically target chemokine receptors involve antagonists that block binding of native chemokines to their receptors and the subsequent receptor signaling responses. By preventing stimulation of downstream effectors that facilitate internalization, antagonists typically stabilize receptors on the cell surface and thus make them available for re-engagement with endogenous agonists upon antagonist dissociation. However, an alternative approach is to minimize the presence of target receptors on the cell surface to render them less accessible to extracellular ligands via “functional antagonism” (19, 24). This strategy was shown to be effective in inhibiting CCR5-mediated HIV entry due to the ability of certain chemokine variants to promote CCR5 sequestration inside the cell (16, 19, 22). However, the mechanisms by which such ligands retain CCR5 inside the cell are poorly understood. To define the ligand-dependent trafficking mechanisms, we employed APEX2 labeling coupled with proteomics (25–27, 55), which allowed for an unbiased identification of proteins in direct contact or close proximity to CCR5 in cells stimulated with WT CCL5 or its sequestering variants, 6P4 and 5P14 (21, 22). By monitoring clusters of labeled proteins based on their cellular compartment localization and the time dependent profiles of the labeling, it was possible to gain insight into the different receptor trafficking patterns induced by the three ligands.

Our results revealed that the three CCL5 ligands exhibit major differences in receptor internalization, subcellular trafficking, recycling and degradation. Striking differences were observed not only in the amount of receptor at various subcellular locations, but also in the receptor trafficking kinetics. 6P4 induced the fastest internalization of CCR5 and allowed receptors to enter recycling pathways as well as late endosomal-lysosomal compartments for degradation. It also directed receptor into Golgi and TGN more so than the other ligands and induced more labeling of proteins from all trafficking pathways, likely by overwhelming multiple trafficking pathways with a high level of internalized receptor **(Figure S7A)**. This mode of CCR5 intracellular sequestration is similar to that previously described for a synthetically modified super-agonist variant PSC-CCL5 (56). By contrast with 6P4 and CCL5, 5P14 induced the slowest receptor internalization and slowest redistribution into other subcellular locations. Nevertheless, it still directed receptors into recycling pathways as well as lysosomal degradation pathways, although it induced less receptor accumulation in the Golgi and TGN, consistent with a prior study (21). Despite inducing a lower level of internalized receptor compared to WT CCL5, 5P14 exhibited a strong preference for late endosome-lysosomal trafficking **(Figure S7A)**.

These observations were supported by bystander BRET assays assessing receptor (CCR5-RlucII) association with different endocytic compartments (early endosomes represented by rGFP-FYVE, recycling endosomes by rGFP-Rab4, and late endosomes by rGFP-Rab7). After normalizing the receptor level in each compartment against the level of internalization, it appears that 6P4 does not preferentially direct CCR5 into any one compartment: they all showed high receptor accumulation **(Figure S7B)**. On the other hand, 5P14 exhibited significantly higher levels of receptor in Rab7-labeled late endosomes relative to the amount of the internalized receptor **(Figure S7B)**. In stark contrast, CCL5 drove receptors into recycling pathways but exhibited negligible degradation and trafficking to the Golgi or TGN. Taken together, it appears that 6P4 sequesters CCR5 by rapidly internalizing a significant amount of receptor and driving it into the Golgi/TGN and along lysosomal degradation pathways, while 5P14 causes slow receptor internalization and recycling as well as induction of receptor degradation.

The trafficking behavior of CCR5 impacts its ability to scavenge chemokine, which is not surprising given the dependence of scavenging on receptor internalization and recycling. CCR5 scavenged CCL5 more efficiently than 6P4 and 5P14: even though 6P4 induces a faster rate and higher degree of CCR5 internalization compared to CCL5, a smaller percentage of receptors return to the cell surface to continuously bind and internalize ligand as required for scavenging. In the case of 5P14, receptor internalization and recycling were slower than that induced by 6P4 and CCL5, consistent with the decreased scavenging of 5P14. By virtue of completely blocking ligand-dependent CCR5 internalization, β-arrestin1/2 knockout fully abrogated scavenging of all ligands by CCR5.

How do these CCL5 ligands trigger the different subcellular trafficking patterns of CCR5? From the proximity labeling data, β-arrestin1/2 emerged as one of the most important master regulators. The level and kinetics of β-arrestin1/2 recruitment to CCR5 following treatment with the different ligands matched the ligand-dependent CCR5 internalization profiles. Knocking out β-arrestin1/2 eliminated ligand-induced internalization for all ligands, also attesting to its dominant role. Finally, numerous clathrin-mediated endocytosis regulators had similar labeling patterns as β-arrestin1/2, and inhibition of CME by pharmacological inhibitors significantly decreased ligand-induced internalization. Thus, the ligand-dependent differences in receptor internalization seem to be defined by β-arrestin1/2 recruitment and subsequent CME.

GRKs were not observed in the proximity labeling data, consistent with the transient nature of their interactions with GPCRs (57). However, receptor phosphorylation is a pre-requisite for β-arrestin1/2 interactions (23, 58) and thus we hypothesized that the level and pattern of GRK phosphorylation may play important roles in ligand-dependent internalization of CCR5. Corroborating this, β-arrestin2 recruitment to CCR5 and CCR5 internalization showed different sensitivities to knockout of different GRK subtypes in a ligand-dependent manner. For CCL5 and 6P4, GRK2/3 played more important roles than GRK5/6 in recruiting β-arrestin2 and driving CCR5 internalization. Of the two ligands, 6P4 showed the strongest effects **(Figure 3A, B)**, likely due to its higher potency and efficacy in inducing G protein activation **(Figure S1B)** which in turn results in stronger phosphorylation by GRK2/3. By contrast, 5P14 treated CCR5 was more sensitive to GRK5/6, presumably because it has a reduced capacity to activate G proteins compared to the other ligands, rendering it less susceptible to phosphorylation by the G protein-dependent kinases GRK2 and GRK3. These ligand-dependent results likely reflect different levels and patterns of “phosphorylation barcodes” (59) by the different GRKs, leading to differences in β-arrestin1/2 interactions and arrestin scaffolded proteins that in turn have consequences on internalization and trafficking. In particular, we identified 1433ζ from our APEX dataset as a protein that may chaperone CCR5 in a [5P14]CCL5-specific manner as a consequence of a GRK5/6 mediated phosphorylation pattern that is sufficient to promote its interaction with CCR5 but not the interaction of CCR5 with β-arrestin1/2.

The precise influence of β-arrestin1/2 on CCR5 subcellular trafficking is more difficult to parse out. However, 6P4 and CCL5 induced persistent CCR5-β-arrestin2 interactions and recruited β-arrestin2 to early endosomes, whereas 5P14 induced more transient interactions. Similar observations were previously made based on immunoprecipitation experiments and speculated to affect CCR5 endocytic trafficking (21). The persistently receptor-bound β-arrestins could negatively regulate CCR5 degradation via direct interaction with the Endosomal Sorting Complex Required for Transport (ESCRT) machinery, as they have been reported to do for CXCR4 (60, 61). Indeed, our APEX proximity labeling data did show that both 6P4 and 5P14, but not WT CCL5, triggered strong labeling of ESCRT components such as STAM1/2 and HGS **(Figure 5A)**. Since 5P14 induced only transient β-arrestin association with CCR5 and did not recruit them to early endosomes, the lack of β-arrestin interactions with ESCRT proteins in endosomal compartments may lead to more receptor degradation through the endo-lysosomal pathway compared to the other ligands **(Figure 5 and Figure S7)**. On the other hand, with stronger and more persistent β-arrestin interactions, 6P4 and CCL5-treated CCR5 were found more in TGN and recycling pathways, respectively. This said, the mechanisms that selectively promoted the proximity to ESCRT proteins of 5P14- and 6P4-stimulated CCR5 remain elusive. In addition, mechanisms independent of β-arrestin2 could also promote 6P4- and 5P14-mediated increases in receptor degradation. We also found increased labeling of components of the alternative ESCRT0 complex such as TOLIP, Endofin (ZFY16) and TOM1 after 6P4 or 5P14 treatment, which targets ubiquitinated proteins for lysosomal degradation. It could be that engineered chemokines trigger specific ubiquitination of CCR5 distinct from the native chemokine CCL5. Other mechanisms may involve prolonged complexation of engineered chemokines with CCR5, potentially in the acidic environment in the endolysosomes, strong subtype G protein signaling bias (20), or other as yet unknown aspects of their unique pharmacology.

While successfully revealing the ligand-dependent intracellular trafficking patterns of CCR5, our study also reveals two important limitations of APEX proximity biotinylation as an experimental technique. First, APEX preferentially labels proteins that are abundant in the cell; as a result, even after removing the bulk of highest abundance proteins prevalent in proximity biotinylation experiments (62), the dataset is dominated by trafficking and housekeeping proteins while low abundance signaling proteins are not well-represented. The use of data-dependent acquisition (DDA) in MS further aggravates the problem. Although one might be able to highlight interesting low-abundance associations by focusing only on proteins with a small number of peptides, this approach is hard to formalize and/or automate for the discovery of direct regulators of CCR5 trafficking and signaling. Comparisons with temporal profiles of bystander/proximity signals from APEX-tagged compartment markers may help to better prioritize such proteins (63).

Additionally, we found that aspects of constitutive receptor trafficking are very hard to deconvolute from APEX proximity biotinylation data. This is because a receptor that constitutively internalizes and recycles is simultaneously present on all endomembranes and labels their resident proteins both in the basal (no agonist) and agonist-stimulated conditions. Agonist stimulation triggers a shift in relative abundance of the receptor at different organelles (e.g. a decrease of the PM population with simultaneous increase in the endosomes), but this still happens in the background of constitutive presence of the receptor in many compartments throughout the cell. Monitoring time-dependent changes in the labeled proximal proteins provides insight into changes between the basal state and the agonist stimulated state, but it does not say much about the basal state itself. Agonist-induced redistribution explains perplexing decreases in CCR5 association with selected compartment-resident proteins (e.g. GOGA2/5). In other words, APEX as well as many other experimental methods (e.g. BRET or IF) are limited in describing the constitutive trafficking of the receptor and instead report agonist-induced changes which gives a false sense of the overall trafficking of a receptor that can be dominated by constitutive processes. The pre-label flow cytometry-based internalization is one of very few methods suitable for characterizing constitutive GPCR trafficking.

In summary, we demonstrated that ligand-dependent internalization of CCR5 is regulated through interactions with β-arrestin1/2 and clathrin-mediated endocytosis. As a master regulator of trafficking, β-arrestin1/2 may also contribute to the post-internalization fate of CCR5 stimulated by some ligands, e.g. by interfacing with ESCRT and other trafficking machinery. However, with other ligands, different mechanisms may be involved. The ligand binding geometry and residence time on the receptor may alter receptor conformation, G protein activation, phosphorylation levels and patterns, and β-arrestin1/2 recruitment levels, as well as kinetics and persistence of receptor complexes with intracellular effectors. Further insight into the ligand-dependent trafficking mechanisms of CCR5 may therefore emerge from deeper investigations of its interactions with other components of the trafficking machinery. In turn, it may be possible to exploit such knowledge for the development of potent CCR5 sequestering agents in the treatment of disease where blocking CCR5 function is desired.

## Materials and Methods

### DNA Constructs

The CCR5 in pcDNA3.1 Hygro(+) was created by PCR amplification of the coding region of N-terminally Flag-tagged CCR5 with subsequent subcloning of the product into pcDNA3.1 Hygro(+) under the CMV (human cytomegalovirus) promoter with the following primers: 5’-CCGGTACCGACTATCAGGTC, 5’- AAGCGGCCGCCTTAAGTCCCACACTGATTTCCTG. For the CCR5-APEX fusion construct, the APEX2 sequence was PCR amplified from pcDNA3-CYTO-APEX2 vector kindly gifted by Dr. Mark von Zastrow (University of California, San Francisco) (25) and subcloned into Flag-CCR5-pcDNA3.1 Hygro+ vector at the C-terminus of CCR5, with a linker (amino acid sequence: AAASGS) separating the two, with the following primers: 5’- ACAAGGACGATGATGACAAACC, 5’- GGTACCGGGTTTGTCATCATC, 5’- CGAGCAGGAAATCAGTGTGGGACTGTCGGGCTCGGGAAAGTC, 5’- CAGTCCCACACTGATTTCCTG, 5’- TAATGACTCGAGTCTAGAGGGCC, 5’- CGGGCCCTCTAGACTCGAGTCATTACGAGCCCGAGGCATC). The spatial references pcDNA3-PM-APEX2, pcDNA3-CYTO-APEX2, and pcDNA3-ENDO-APEX2 were also kindly gifted by Dr. Mark von Zastrow (University of California, San Francisco) (25). The pcDNA3.1(+) constructs of β-arrestin1-RlucII, β-arrestin2-RlucII, rGFP-CAAX, rGFP-Rab4 were kindly gifted by Dr. Michel Bouvier (Université de Montréal, Canada) (36). The rGFP-Rab7 construct was generated by PCR amplification of the Rab7 coding sequence of EGFP-Rab7A, a gift from Qing Zhong (UC Berkeley, CA, USA) and inserted in-frame into the XbaI and PmeI sites of rGFP-Rab4 as previously reported (31). The CCR5-RlucII construct was created by PCR amplification of the coding region of CCR5; products were subcloned in-frame at the N-terminus of the RlucII sequence into the pcDNA3.1 RlucII vector with the following primers: 5’-CCGGTACCGACTATCAGGTC, 5’-CGCGGATCCCCTGGTTCCAGTCCCACACTGATTTCCTG. To create the FLAG-SNAP-CCR5 construct, the coding sequence of a pcDNA3.1(+) CCR5 vector was PCR amplified and inserted into a pRK5–FLAG-ST-CXCR4 plasmid (64) with the following primers: 5’- CGCACGCGTGACTATCAGGTCAGCTCCC, 5’- ACGGGCCCTCATTACAGTCCCACACTGATTTCC, containing the mGlu5 receptor signal peptide that promotes proper receptor trafficking to the cell surface (kind gift of Dr. Angélique Levoye, University of Paris, France).

### Cell lines

All KO and parental (WT) control HEK293 cells were a kind gift of Dr. Asuka Inoue (Tohoku University, Japan). A dual β-arrestin1 and β-arrestin2 knockout (β-arrestin1/2 KO) was prepared by CRISPR/Cas9 targeting of ARRB1 and ARRB2 as described previously (65). Similarly, CRISPR/Cas9 was used to generate GRK2/3 KO, GRK5/6 KO and GRK2/3/5/6 KO in the HEK293A cell line (66). Stable chemokine receptor-expressing cells (WT CCR5 and CCR5-APEX) and spatial reference-expressing cells were generated by transfecting HEK293T with expression vectors using the TransIT-LT1 transfection reagent (Mirus Bio, MIR2305). Stable transfectants were subsequently selected using 100 ug/mL Hygromycin B (Life Technologies, 10687010) for chemokine receptor-expressing cells or 500 ug/mL G418 for spatial reference cells. Receptor surface expression was confirmed by flow cytometry (Guava EasyCyte™ 8HT, Luminex) with an anti-hCCR5-APC (Allophycocyanin) conjugated antibody (eBioscience, 501123098). Cell lines were cultured in Dulbecco’s modified Eagle’s medium (DMEM) supplemented with GlutaMax (Gibco), 10% fetal bovine serum (FBS), and selecting reagent, and grown at 37°C with 5% CO_2_.

### APEX proximity labeling and streptavidin pull down

CCR5-APEX stable cells were cultured in 15 cm dishes in DMEM with GlutaMax (Gibco) and 10% fetal bovine serum (FBS) until confluent. The cells were then pre-incubated with 500 μM biotinyl tyramide (Sigma, SML2135) for 30 min at 37°C and then treated with 100 nM corresponding CCL5 ligands for the indicated periods of time. Labeling was initiated with 1 mM H_2_O_2_ for 45 seconds, and then quenched with ice cold quenching buffer composed of 1 mM CaCl_2_, 10 mM sodium ascorbate, 1 mM Trolox and 1 mM sodium azide in PBS. The cells were harvested and lysed in RIPA buffer containing 10 mM sodium ascorbate, 1 mM Trolox, 1 mM sodium azide, 1mM DTT, protease inhibitor (Sigma, 5056489001) and Halt phosphatase inhibitor (Thermo Fisher Scientific, PI78420). After clarifying the lysates by centrifugation at 17,000 x g, at 4 °C for 15 min, the lysates were incubated with streptavidin magnetic beads (Thermo Fisher Scientific, 88817) with rotation at 4°C overnight. The beads were then washed three times with buffer containing 4 M urea, 0.5% DDM (w/v), 100 mM sodium phosphate pH 8, followed by three washes with 4M urea, 100mM sodium phosphate pH 8 buffer, and finally additional wash with 50 mM HEPES, pH 8.5. The beads were resuspended in 50 mM HEPES pH 8.5 and stored at - 80°C before further processing.

### Quantitative Multiplex Proteomics

Please see Supporting Information, Extended Materials and Methods for details.

### Enhanced bystander bioluminescence resonance energy transfer (ebBRET) assay

HEK293 cells were seeded into 6-well plates (750,000 cells per well) and transfected the next day using Mirus TransIT-LT1 transfection reagent at ∼70% confluency. Cells were transfected with a BRET donor (32.5 ng of Receptor-RlucII or 19.8 ng of β-arrestin1- or β-arrestin2-RlucII per well) along with 130 ng of BRET acceptor per well (for example, rGFP-CAAX, rGFP-Rab4, rGFP-Rab11 or rGFP-Rab7). All assays were performed ∼30 h after transfection, using previously described methods (31) with minor modifications. Specifically, transfected cells were detached with 0.25ml/well Accutase (Invitrogen^TM^, 00-4555-56) for 3 min at 25°C, lifted from the 6-well plates by pipetting, and seeded into a 96-well white microplate (Tecan, 30122300) at 100,000 cells per well in pre-warmed Tyrode’s buffer (140 mM NaCl, 2.7 mM KCl, 1 mM CaCl_2_, 12 mM NaHCO_3_, 5.6 mM D-glucose, 0.5 mM MgCl_2_, 0.37 mM NaH_2_PO_4_, 25 mM HEPES, pH 7.4). A cell permeable RlucII substrate, Prolume Purple (NanoLight Technologies) was added to the wells at a final concentration of 5 μM, ∼3 to 6 min before BRET measurements. Three baseline BRET measurements were performed approximately 1 min apart, followed by the addition of the indicated concentrations of chemokines. Subsequent BRET measurements were taken approximately 1 min apart for 50 min or the indicated times.

For evaluation of receptor recycling, cells were transfected and seeded as described above. Following 30 min of chemokine stimulation and BRET measurements approximately 2 min apart, inhibitor Maraviroc (Sigma, PZ0002) was added to the cells at a final concentration of 5 μM and subsequent BRET measurements were taken for an additional 40 min.

All BRET measurements were read using a VictorX Light luminescence reader (Perkin Elmer, Waltham, MA, USA) or Spark microplate reader (Tecan, Männedorf, Switzerland). Values are presented as the fold change over mock-treated.

### Flow cytometry-based internalization assay

For the **pre-label assay**, HEK293 cells stably expressing CCR5 were labeled with 2ug/ml rabbit anti-Flag tag antibody (F7425, Sigma, St. Louis, MO, USA) in FACS buffer (PBS with 0.5% bovine serum albumin (BSA)) for 30 min on ice and protected from light.

Unbound antibody was removed by repeated washes with FACS buffer. Cells were then resuspended in Assay Buffer (DMEM, 0.5% BSA) containing 100 nM CCL5 ligands (CCL5, 6P4, or 5P14) or without ligands, and then either held at 4°C (which prevents receptor trafficking and internalization) or transferred to 37°C (which allows for normal receptor trafficking) and incubated for 1 h. After incubation, cells were transferred to wet ice and the remaining surface receptor was labeled with 1:100 anti-rabbit antibody conjugated to PE (F0110, R&D Systems, Minneapolis, MN, USA) in FACS buffer for 30 min on ice and protected from light. Following 3 washes with FACS buffer, CCR5 expression was assessed by flow cytometry using a Guava EasyCyte™ 8HT flow cytometer (Luminex). Data was analyzed with FlowJo software (FlowJo, Ashland, OR, USA). The geometric mean fluorescent intensity of analyzed cells was used to quantify surface expression of CCR5 and compared to non-internalized control to determine relative percent of receptor remaining at the surface. For the **post-label assay,** all steps are the same as the pre-label assay except that the primary antibody staining was performed <u>after</u> the ligand incubation.

### Receptor degradation assay

HEK293 cells were seeded in 10 cm dishes and transfected at 70-80% confluency with FLAG-CCR5 using Mirus TransIT-Lt1 transfection reagent. After 24 h, the cells were re-plated into a 6-well plate, 3.5x10^5^/ml cells per well in 2.5ml DMEM media. After 48 h, cells were pre-treated with 50 μg/mL cycloheximide for 15 min at 37°C and incubated in the same media with or without 100 nM chemokine ligands for 4 h at 37°C. Following incubation, cells were placed on ice, rinsed with PBS, and lysed in RIPA lysis buffer (Thermo Fisher Scientific) containing cOmplete™ Mini EDTA-free Protease Inhibitor Cocktail (Roche). Cell lysates were collected and rotated end-over-end for 1 h at 4°C. Total protein concentrations were determined by Pierce™ Rapid Gold bicinchoninic acid (BCA) assay (Thermo Fisher Scientific). Equivalent amounts of lysates were resolved by SDS-PAGE and transferred to a nitrocellulose membrane (Bio-Rad). Membranes were blocked with 5% nonfat dry milk (Bio-Rad) diluted in wash buffer (50 mm Tris-HCl, pH 7.4, 150 mM NaCl, 0.1% Tween 20) and subsequently washed in the same buffer. Blots were incubated overnight at 4°C with 1:1000 anti-FLAG rabbit antibody (F4725, Millipore Sigma) and 1:1000 anti-Actin mouse antibody (A3853, Millipore Sigma) diluted in wash buffer containing 5% BSA. Membranes were washed and probed with 1:10000 of the corresponding secondary IRDye® 800CW antibody and 680RD antibody (LI-COR Biosciences) diluted in wash buffer containing 5% BSA at room temperature for 1 h. After final washes, membranes were imaged on a Chemidoc^TM^ MP imaging system (BioRad). Densitometry was performed using Chemidoc^TM^ Image Lab software (BioRad).

### Confocal microscopy

HEK293 cells expressing Flag-SNAP-CCR5 were seeded onto 10-mm glass-bottom dishes (FluoroDish, FD3510, WPI) previously coated with 5 μg/ml fibronectin. The next day, cells were labeled with 5 μM cell-impermeable SNAP-Surface Alexa Fluor 488 (New England Biolabs) in complete media (DMEM + 10% FBS) on ice in the dark. After 1 h, the cells were washed with ice-cold complete media and fixed with 4 % paraformaldehyde after being incubated at 37°C with 100 nM of different agonists for 1 h. Control cells were fixed directly after labeling. Fixed cells were permeabilized using PBS containing 0.1% saponin for 30 min and blocked using PBS with 1% BSA and 1% goat serum for 1 h before labeling with 1:100 dilution of rabbit polyclonal anti-TGN-46 antibody (Novus biologicals, NBP1-49643). After primary antibody labeling, cells were washed using PBS containing 0.1% saponin and incubated using goat anti-Rabbit IgG Alexa Fluor™ Plus 647 (1:1000 dilution) for 1 h. Cells were washed and incubated briefly with Hoechst 33342 before imaging using an Eclipse Ti2-E (Nikon) equipped with a CSU-X1 (Yokogawa) spinning disk field scanning confocal system. Image analysis was performed using ImageJ (NIH, Bethesda, MD, USA) software and colocalization was measured using the JaCoP plugin.

### Scavenging assay

Non-receptor-expressing or stable CCR5-expressing HEK293 cells and corresponding β-arrestins KO cells were seeded in triplicate in 96-well dishes at 50,000 cells/well in DMEM/10%FBS media and allowed to adhere for ∼8 h. Subsequently, media was replaced with DMEM/10% FBS media containing 5 nM CCL5 ligands (CCL5, 6P4 or 5P14) and 10ug/ml heparin, and incubated for ∼16 h. Remaining chemokine levels in the supernatants of cultured cells were measured in triplicate using the commercially available colorimetric LEGEND MAX^TM^ human CCL5 ELISA kit (BioLegend, 440807) according to the manufacturer’s instructions and read with a SpectraMax M5 plate reader (Molecular Devices, Sunnyvale, CA, USA). Remaining levels of exogenous chemokine are reported as the percentage of levels in supernatants of the corresponding non-receptor expressing cells.

### Structure modeling

3D models of complexes between C-terminally Ser-phosphorylated CCR5 with [5P14]CCL5 and dimeric 1433ζ were built from amino acid sequences of CCR5(aa 1-352), [5P14]CCL5 (aa 1-69), and two copies of 1433ζ (aa 1-245) using AlphaFold3 (53) through the AlphaFold Server (https://golgi.sandbox.google.com/). A single phosphoserine modification was applied on S336, S337, S342 or S349. 5 models were built per complex. Models were evaluated by the predicted per-residue pLDDT scores and inspected visually. The model with the highest pLDDT score is shown in **Fig. 8**.

### Statistics

All data points are the mean ± SE of three independent experiments conducted separately (biological replicates). Within each independent experiment there are three technical replicates for each condition conducted in parallel, except for the APEX proximity labeling and degradation assays which only have biological replicates. All data were analyzed using GraphPad Prism with statistically significant differences (P < 0.05) using unpaired t test with Welch’s correction. For BRET internalization, β-arrestin recruitment, and recycling, the area under the curve (AUC) was determined using GraphPad Prism and statistical significance was determined using the test mentioned above. For microscopy experiments, more than 200 Z-stack images containing > 30 cells per condition were analyzed for Pearson’s Correlation Coefficient (PCC) with statistically significant differences (P<0.0001) determined using the Mann–Whitney test.

## Acknowledgments

T.M.H and I.K. acknowledge support from NIH (R01 AI161880, R01 GM136202, R01 CA254402). I.K. acknowledges additional support from NIH (R21 AI149369, R21 AI156662). S.G. was partly supported by the Cancer Research Institute Irvington Postdoctoral Fellowship. D.J.G. and S.M. were supported by the Collaborative Center for Multiplexed Proteomics at the UCSD School of Medicine.

## References

1. C. A. Flanagan, Advances in Pharmacology. Adv Pharmacol San Diego Calif 70, 215–263 (2014).

2. A. Brelot, L. A. Chakrabarti, CCR5 revisited: how mechanisms of HIV entry govern AIDS pathogenesis. J Mol Biol 430, 2557–2589 (2018).

3. P. M. Murphy, Viral exploitation and subversion of the immune system through chemokine mimicry. Nat Immunol 2, 116–122 (2001).

4. P. E. Sax, FDA approval: maraviroc. AIDS Clin Care 19, 75 (2007).

5. M. T. Joy et al., CCR5 Is a Therapeutic Target for Recovery after Stroke and Traumatic Brain Injury. Cell 176, 1143–1157.e1113 (2019).

6. S. Lefere, L. Devisscher, F. Tacke, Targeting CCR2/5 in the treatment of nonalcoholic steatohepatitis (NASH) and fibrosis: opportunities and challenges. Expert Opinion on Investigational Drugs 29, 89–92 (2020).

7. L. A. Boven, L. Montagne, H. S. Nottet, C. J. De Groot, Macrophage inflammatory protein-1alpha (MIP-1alpha), MIP-1beta, and RANTES mRNA semiquantification and protein expression in active demyelinating multiple sclerosis (MS) lesions. Clin Exp Immunol 122, 257–263 (2000).

8. A. Zernecke et al., Deficiency in CCR5 but not CCR1 protects against neointima formation in atherosclerosis-prone mice: involvement of IL-10. Blood 107, 4240–4243 (2006).

9. M. N. Ajuebor, C. M. Hogaboam, S. L. Kunkel, A. E. Proudfoot, J. L. Wallace, The chemokine RANTES is a crucial mediator of the progression from acute to chronic colitis in the rat. J Immunol 166, 552–558 (2001).

10. C. E. C. d. Oliveira et al., CC Chemokine Receptor 5: The Interface of Host Immunity and Cancer. Dis Markers 2014, 126954 (2014).

11. L. Vangelista, S. Vento, The Expanding Therapeutic Perspective of CCR5 Blockade. Frontiers in Immunology 8, 1981 (2018).

12. J. Zhao et al., Chemokine receptor CCR5 functionally couples to inhibitory G proteins and undergoes desensitization. Journal of Cellular Biochemistry 71, 36–45 (1998).

13. I. Aramori et al., Molecular mechanism of desensitization of the chemokine receptor CCR-5: receptor signaling and internalization are dissociable from its role as an HIV-1 co-receptor. The EMBO Journal 16, 4606–4616 (1997).

14. B. S. Christmann, J. M. Moran, J. A. McGraw, R. M. Buller, J. A. Corbett, Ccr5 regulates inflammatory gene expression in response to encephalomyocarditis virus infection. Am J Pathol 179, 2941–2951 (2011).

15. M. Wong et al., Rantes activates Jak2 and Jak3 to regulate engagement of multiple signaling pathways in T cells. J Biol Chem 276, 11427–11431 (2001).

16. M. Mack et al., Aminooxypentane-RANTES induces CCR5 internalization but inhibits recycling: a novel inhibitory mechanism of HIV infectivity. J Exp Med 187, 1215–1224 (1998).

17. M. Oppermann, Chemokine receptor CCR5: insights into structure, function, and regulation. Cell Signal 16, 1201–1210 (2004).

18. F. Hüttenrauch, B. Pollok-Kopp, M. Oppermann, G protein-coupled receptor kinases promote phosphorylation and beta-arrestin-mediated internalization of CCR5 homo- and hetero-oligomers. J Biol Chem 280, 37503–37515 (2005).

19. G. Simmons et al., Potent Inhibition of HIV-1 Infectivity in Macrophages and Lymphocytes by a Novel CCR5 Antagonist. Science 276, 276–279 (1997).

20. E. Lorenzen et al., G protein subtype–specific signaling bias in a series of CCR5 chemokine analogs. Sci Signal 11, eaao6152 (2018).

21. C. Bönsch, M. Munteanu, I. Rossitto-Borlat, A. Fürstenberg, O. Hartley, Potent Anti-HIV Chemokine Analogs Direct Post-Endocytic Sorting of CCR5. Plos One 10, e0125396 (2015).

22. H. Gaertner et al., Highly potent, fully recombinant anti-HIV chemokines: Reengineering a low-cost microbicide. Proc National Acad Sci 105, 17706–17711 (2008).

23. E. Martins et al., Arrestin Recruitment to C-C Chemokine Receptor 5: Potent C-C Chemokine Ligand 5 Analogs Reveal Differences in Dependence on Receptor Phosphorylation and Isoform-Specific Recruitment Bias. Mol Pharmacol 98, 599–611 (2020).

24. K. LaMontagne et al., Antagonism of Sphingosine-1-Phosphate Receptors by FTY720 Inhibits Angiogenesis and Tumor Vascularization. Cancer Research 66, 221–231 (2006).

25. B. T. Lobingier et al., An Approach to Spatiotemporally Resolve Protein Interaction Networks in Living Cells. Cell 169, 350–360.e312 (2017).

26. J. Paek et al., Multidimensional Tracking of GPCR Signaling via Peroxidase-Catalyzed Proximity Labeling. Cell 169 (2017).

27. B. J. Polacco et al., Profiling the diversity of agonist-selective effects on the proximal proteome environment of G protein-coupled receptors. Biorxiv 10.1101/2022.03.28.486115, 2022.2003.2028.486115 (2022).

28. J.-E. Turner et al., Protective role for CCR5 in murine lupus nephritis. American Journal of Physiology-Renal Physiology 302, F1503–F1515 (2012).

29. A. E. Cardona et al., Scavenging roles of chemokine receptors: chemokine receptor deficiency is associated with increased levels of ligand in circulation and tissues. Blood 112, 256–263 (2008).

30. C. T. Gilliland, C. L. Salanga, T. Kawamura, J. Trejo, T. M. Handel, The Chemokine Receptor CCR1 Is Constitutively Active, Which Leads to G Protein-independent, β-Arrestin-mediated Internalization. J Biol Chem 288, 32194–32210 (2013).

31. T. M. Shroka, I. Kufareva, C. L. Salanga, T. M. Handel, The dual-function chemokine receptor CCR2 drives migration and chemokine scavenging through distinct mechanisms. Sci Signal 16, eabo4314 (2023).

32. A. Thompson et al., Tandem Mass Tags: A Novel Quantification Strategy for Comparative Analysis of Complex Protein Mixtures by MS/MS. Anal Chem 75, 1895–1904 (2003).

33. M. E. Ritchie et al., limma powers differential expression analyses for RNA-sequencing and microarray studies. Nucleic Acids Res 43, e47 (2015).

34. M. Ashburner et al., Gene Ontology: tool for the unification of biology. Nature Genetics 25, 25–29 (2000).

35. J. A. Paulo et al., Quantitative mass spectrometry-based multiplexing compares the abundance of 5000 S. cerevisiae proteins across 10 carbon sources. J Proteomics 148, 85–93 (2016).

36. Y. Namkung et al., Monitoring G protein-coupled receptor and β-arrestin trafficking in live cells using enhanced bystander BRET. Nat Commun 7, 12178 (2016).

37. T. Ferain et al., Agonist-Induced Internalization of CC Chemokine Receptor 5 as a Mechanism to Inhibit HIV Replication. J Pharmacol Exp Ther 337, 655–662 (2011).

38. M. H. Dreyling et al., The t(10;11)(p13;q14) in the U937 cell line results in the fusion of the AF10 gene and CALM, encoding a new member of the AP-3 clathrin assembly protein family. Proc Natl Acad Sci U S A 93, 4804–4809 (1996).

39. A. Benmerah et al., The tyrosine kinase substrate eps15 is constitutively associated with the plasma membrane adaptor AP-2. J Cell Biol 131, 1831–1838 (1995).

40. D. Dutta, C. D. Williamson, N. B. Cole, J. G. Donaldson, Pitstop 2 is a potent inhibitor of clathrin-independent endocytosis. Plos One 7, e45799 (2012).

41. A. McCluskey et al., Building a better dynasore: the dyngo compounds potently inhibit dynamin and endocytosis. Traffic 14, 1272–1289 (2013).

42. S. S. Ferguson et al., Role of beta-arrestin in mediating agonist-promoted G protein-coupled receptor internalization. Science 271, 363–366 (1996).

43. O. B. Goodman, Jr., J. G. Krupnick, V. V. Gurevich, J. L. Benovic, J. H. Keen, Arrestin/clathrin interaction. Localization of the arrestin binding locus to the clathrin terminal domain. J Biol Chem 272, 15017–15022 (1997).

44. J. S. Paradis et al., Receptor sequestration in response to β-arrestin-2 phosphorylation by ERK1/2 governs steady-state levels of GPCR cell-surface expression. Proc National Acad Sci 112, E5160–E5168 (2015).

45. T. J. Cahill, 3rd et al., Distinct conformations of GPCR-β-arrestin complexes mediate desensitization, signaling, and endocytosis. Proc Natl Acad Sci U S A 114, 2562–2567 (2017).

46. Z. Yang et al., Phosphorylation of G protein-coupled receptors: from the barcode hypothesis to the flute model. Mol Pharmacol 92, mol.116.107839 (2017).

47. C. Mauvezin, T. P. Neufeld, Bafilomycin A1 disrupts autophagic flux by inhibiting both V-ATPase-dependent acidification and Ca-P60A/SERCA-dependent autophagosome-lysosome fusion. Autophagy 11, 1437–1438 (2015).

48. A. Ariel et al., Apoptotic neutrophils and T cells sequester chemokines during immune response resolution through modulation of CCR5 expression. Nat Immunol 7, 1209–1216 (2006).

49. V. Obsilova, T. Obsil, Structural insights into the functional roles of 14-3-3 proteins. Front Mol Biosci 9, 1016071 (2022).

50. M. J. van Hemert, H. Y. Steensma, G. P. H. van Heusden, 14-3-3 proteins: key regulators of cell division, signalling and apoptosis. BioEssays 23, 936–946 (2001).

51. L. Yuan et al., 14-3-3 signal adaptor and scaffold proteins mediate GPCR trafficking. Sci Rep-uk 9, 11156 (2019).

52. F. Madeira et al., 14-3-3-Pred: improved methods to predict 14-3-3-binding phosphopeptides. Bioinformatics 31, 2276–2283 (2015).

53. J. Abramson et al., Accurate structure prediction of biomolecular interactions with AlphaFold 3. Nature 630, 493–500 (2024).

54. M. Oppermann, M. Mack, A. E. Proudfoot, H. Olbrich, Differential effects of CC chemokines on CC chemokine receptor 5 (CCR5) phosphorylation and identification of phosphorylation sites on the CCR5 carboxyl terminus. J Biol Chem 274, 8875–8885 (1999).

55. C. D. Go et al., A proximity biotinylation map of a human cell. Biorxiv 10.1101/796391, 796391 (2019).

56. J. M. Escola, G. Kuenzi, H. Gaertner, M. Foti, O. Hartley, CC chemokine receptor 5 (CCR5) desensitization: cycling receptors accumulate in the trans-Golgi network. J Biol Chem 285, 41772–41780 (2010).

57. Y. He et al., Molecular assembly of rhodopsin with G protein-coupled receptor kinases. Cell Res 27, 728–747 (2017).

58. P. Isaikina et al., A key GPCR phosphorylation motif discovered in arrestin2⋅CCR5 phosphopeptide complexes. Mol Cell 83, 2108--2121.e2107 (2023).

59. S. B. Liggett, Phosphorylation Barcoding as a Mechanism of Directing GPCR Signaling. Sci Signal 4, pe36–pe36 (2011).

60. R. Malik, A. Marchese, Arrestin-2 interacts with the endosomal sorting complex required for transport machinery to modulate endosomal sorting of CXCR4. Mol Biol Cell 21, 2529–2541 (2010).

61. O. Alekhina, A. Marchese, β-Arrestin1 and Signal-transducing Adaptor Molecule 1 (STAM1) Cooperate to Promote Focal Adhesion Kinase Autophosphorylation and Chemotaxis via the Chemokine Receptor CXCR4. J Biol Chem 291, 26083–26097 (2016).

62. D. Mellacheruvu et al., The CRAPome: a contaminant repository for affinity purification– mass spectrometry data. Nature Methods 10, 730–736 (2013).

63. B. J. Polacco et al., Profiling the proximal proteome of the mu opioid receptor identifies novel regulators of receptor signaling and trafficking. Biorxiv 10.1101/2022.03.28.486115, 2022.2003.2028.486115 (2023).

64. A. Levoye et al., A Broad G Protein-Coupled Receptor Internalization Assay that Combines SNAP-Tag Labeling, Diffusion-Enhanced Resonance Energy Transfer, and a Highly Emissive Terbium Cryptate. Front Endocrinol (Lausanne) 6, 167 (2015).

65. M. O’Hayre et al., Genetic evidence that β-arrestins are dispensable for the initiation of β(2)-adrenergic receptor signaling to ERK. Sci Signal 10 (2017).

66. S. Pandey et al., Intrinsic bias at non-canonical, β-arrestin-coupled seven transmembrane receptors. Mol Cell 81, 4605–4621.e4611 (2021).

67. C. Petosa et al., 14-3-3ζ Binds a Phosphorylated Raf Peptide and an Unphosphorylated Peptide via Its Conserved Amphipathic Groove*. J Biol Chem 273, 16305–16310 (1998).

