## Supplemental Information for "Ligand-Dependent Mechanisms of CC Chemokine Receptor 5 (CCR5) Trafficking Revealed by APEX2 Proximity Labeling Proteomics"

**Supporting Information for**
### CC Chemokine Receptor 5 (CCR5) Ligand-Dependent Trafficking Mechanisms Guided by APEX2 Proximity Labeling Proteomics

Siyi Gu, Svetlana Maurya, Alexis Lona, Leire Borrega-Roman, Catherina Salanga, David Gonzalez, Irina Kufareva, Tracy M. Handel

Skaggs School of Pharmacy and Pharmaceutical Sciences, University of California San Diego, La Jolla, CA, 92093, USA

Department of Pharmacology, University of California San Diego, La Jolla, CA, 92093, USA

* Tracy M. Handel


* Irina Kufareva


#### Extended Materials and Methods.

***Chemokine production.*** CCL5 sequences was cloned into the NdeI/XhoI site of a pET21-based vector with an N-terminal His8 tag followed by an enterokinase recognition site. The desired mutations were introduced into the WT sequence using QuikChange site-directed mutagenesis (Stratagene). For protein production, the corresponding CCL5 plasmid was transformed into E. coli BL21(DE3) cells. Cells were grown at 37 °C in Luria-Bertani medium, and protein expression was induced with isopropyl β-d-thiogalactopyranoside at midlog phase (A600 = 0.6) for 3–4 h. Cells were harvested by centrifugation, resuspended (50 mM Tris, pH 7.5 with 150 mM NaCl) and frozen at −80 °C. Cell pellets were lysed by sonication; inclusion bodies containing CCL5 were collected by centrifugation and dissolved in denaturant (6 M guanidinium chloride, 50 mM NaCl, 50 mM Tris, pH 8.0) and then purified using nickel-nitrilotriacetic acid resin (Qiagen). Refolding of CCL5 variants was done by adding 4mM DTT to the eluant at pH=7-7.5 for 1 hr, and then exchanging the eluant into non-denaturing buffer (50 mM Tris, pH 7.5, 500 mM L-arginine, 1 mM EDTA, 1mM GSSG) and allowing the refolding to proceed overnight at 4 °C. The refolded product was then dialyzed into buffer containing 50mM MES pH 5.7, 50mM NaCl, and treated with enterokinase at 37 °C for 12-18 hrs. The cleaved product was purified once again using nickel-nitrilotriacetic acid resin, followed by reverse phase HPLC on a C18 column, frozen, lyophilized, and stored at −80 °C.

##### Quantitative Multiplex Proteomics

***Sample preparation.*** Biotinylated samples from APEX proximity labeling and streptavidin pull down were analyzed by quantitative multiplex proteomics at the UCSD Collaborative Center for Multiplexed Proteomics (1). Biotinylated proteins bound to magnetic streptavidin beads (Thermo Fisher Scientific, 88817) were precipitated using a magnetic stand (Permagen, MSR6X15) and resuspended in 2mL of 4M Urea in 50mM HEPES. Disulfide bonds were reduced in 5 mM dithiothreitol (DTT) at 47°C for 30 min and free cysteines were alkylated in 15 mM iodoacetamide at room temperature in the dark for 20 min. The alkylation reaction was quenched for 15 min at room temperature with 15mM DTT. Beads with samples were then pulled down using a magnetic stand and washed with 3mL of 1M Urea in 50mM HEPES. Samples were then resuspended in 2mL of 1M urea in 50mM HEPES. Proteins bound to streptavidin beads were digested with 20ug of trypsin (Promega, V5113) at 47°C for 5 hours. Streptavidin beads were pulled down once more using a magnetic stand and the in-solution digest was transferred to a separate Protein LoBind microcentrifuge tube and acidified using trifluoroacetic acid to a final concentration of 1%. Samples were desalted using C18 columns (Waters) using manufacturer protocols and dried under vacuum. Samples were subjected to peptide quantification using a Pierce Colorimetric Peptide Quantification Assay kit per the manufacturer's instructions. 50 µg of each peptide sample was aliquoted for TMT labeling.

***Tandem mass tag labeling.*** The peptide samples were resuspended in a solution of 30% anhydrous acetonitrile (ACN) with 200 mM HEPES, pH = 8.5. Tandem mass tag (TMTpro) labels (Thermo Fisher Scientific, A44520, XC342532) were suspended in anhydrous ACN to a final concentration of 20 mg/mL, and 7 µL added to each resuspended sample. Samples were randomized in the TMT 16plex, with the TMT126 channel used as a bridge. The labeling reaction was allowed to proceed for 1 h at room temperature, after which excess label was quenched by the addition of 8 µL of 5% hydroxylamine for 15 min. 50 µL of 1% (TFA) was added to each sample; samples were then pooled together into their respective TMT-plex and desalted on C18 columns (Waters) as above. The multiplexed samples were dried under vacuum.

***Mass spectrometry-based proteomic analysis.*** Labeled peptides were resuspended in 5% Formic Acid/5% ACN to a final concentration of 1ug/uL for mass spectrometry analysis. All proteome mass spectrometry data were collected on an Orbitrap Fusion mass spectrometer (Thermo Fisher Scientific) with an in line Easy-nLC$^{TM}$ 1200 HPLC system (Thermo Fisher Scientific, LC140). Previously described methods were utilized for data collection (1). Briefly, samples were loaded on a 30cm glass capillary column packed with 0.5cm of 5um C4 resin followed by 0.5cm of 3um C18 resin, and then 29cm of 1.8uM C18 resin. Following sample loading, peptides were eluted using a gradient of 3% ACN/0.125% formic acid (A) and 100% ACN/0.125% formic acid (B) starting at 6% of mixture A and ending in 100% mixture B over a 180 min gradient at a flow rate of 300nL/min, with the column heated to 60°C.

***Data processing and normalization*** Raw mass spectrometry files were searched using the SEQUEST algorithm in Proteome Discoverer 2.5 against the reference proteome for *Homo sapiens* retrieved July 5$^{th}$, 2022 from UniProt server. Data were searched using a precursor mass tolerance of 50 ppm and fragment mass tolerance of 0.6. Static modifications were specified as follows: TMTpro on lysine and N-termini and carbamidomethylation of cysteines. Dynamic modifications were specified to include oxidation of methionine. Resulting peptide spectral matches were filtered at a 0.01 false discovery rate by the Percolator module against the decoy database.

**Bioinformatics analysis**

***Data Availability.*** Proteome data was uploaded to massive.ucsd.edu and the ProteomeXchange Consortium (https://www.proteomexchange.org/). The data can be accessed using the following identifiers: MassIVE MSV000093226 (massive.ucsd.edu) and PXD046588 (ProteomeXchange).

***Code Availability.*** R scripts and code are available at https://github.com/Kufalab-UCSD/CCR5_APEX for project-specific, post-acquisition data processing and https://github.com/Kufalab-UCSD/ProteomicsToolkit for inhouse developed proteomic analysis tools for R coding environments.

***Peptide-spectrum match quant analysis.*** Mass spectrometry peptide spectrum matches (PSMs) were summed to peptides, transformed into natural log scale, and median-centered **(Figure S3A)**. Peptides were remapped to UniProtKB release 2021-04; any peptides that could not be unambiguously assigned to a single protein formed a protein group that was treated separately from constituent proteins. Technical variation (a.k.a. "batch effects") between the TMT 16-plexes was removed using a combined strategy of subtracting batch-aggregate "bridge samples" and linear modeling as implemented in the *limma* R library (2, 3). Quality of batch effect removal was controlled for by comparing adjusted Rand indices between the sample-to-sample correlation dendrogram cut at different heights and the experimental variables **(Figure S3B)** (4). Due to a technical issue, all three samples stimulated by 5P14 for 10 min had a substantially broader quant distribution **(Figure S3C);** because of this, they were omitted from any overall receptor behavior analyses.

Proteins were assigned numerical scores reflecting their likelihood of being experimental contaminants: these scores were calculated based on REPRINT's Contaminant Repository for Affinity Purification (CRAPome, H. sapiens - Proximity Dependent Biotinylation (5); a higher CRAPome score means a greater likelihood of being a contaminant), their quants in our negative control (non-$H_2O_2$-treated) samples (a higher quant in the absence of $H_2O_2$ means a greater likelihood of being a contaminant), and their membership in the Reactome terms "Signal Transduction" or "Vesicle-Mediated Transport" (membership means lower likelihood of being a contaminant). For each protein, random constituent peptides were removed, with the fraction of removed peptides proportional to the protein's contaminant likelihood score. Altogether, the contaminant removal process reduced the number of peptides in the dataset by ~40% but the number of unique proteins only by ~ 10%.

For each peptide in the filtered dataset, statistical significance of variation across all ligand/time conditions was assessed using moderated ANOVA with Benjamin-Hochberg (BH) adjustment for multiple comparisons, as implemented in *limma*. Next, peptide quants were projected onto the proteins using the philosophy of EBProt (6, 7): for this, quants from up to 10 most significant constituent peptides from each protein were made comparable to each other via linear modeling-based batch effect removal and treated as replicate measurements on the protein. This approach ensures that proteins are only prioritized if the patterns of MS quant variations are consistent across all of their constituent peptides. Afterwards, two-way moderated ANOVA with a BH adjustment was calculated on all proteins across all ligand conditions, resulting in the identification of 491 significantly varied proteins ($p < 0.01$).

For **Fig. 1A**, over-representation of Gene Ontology Biological Process (BP) and Cellular Compartment (CC) terms was analyzed in clusterProfiler R library (8): 491 proteins were annotated against a background of 3187 total proteins. Significances of overrepresented terms were adjusted using BH correction for multiple comparisons.

For **Fig. 1B**, pairwise similarity between biotinylation response profiles of different proteins was calculated using the extra sum-of-squares F test (9); p-values were transformed to negative log10 scale (so that larger numbers corresponded to more distinct profiles) and used as protein-protein response profile distances. The resulting distance matrix was used to hierarchically cluster proteins according to their variation profiles across all ligand conditions. A cutoff height of 8 was used to separate proteins into 66 clusters. Clusters were fuzzified via calculating F test-based distances from each dataset protein to each cluster center, and the resulting cluster memberships were combined with proteins' own significance of variation to get membership-corrected significance values. Over-representation of Gene Ontology CC terms within these clusters was analyzed in clusterProfiler as for **Fig. 1A**; significantly varied proteins (membership-corrected $p < 0.05$) within a cluster were compared to the total experiment background.

For **Fig. 8A** and **Fig S8A**, the significance of the difference between the proximity response profiles to [5P14]CCL5 and WT CCL5 was calculated using an F test and plotted against the significance of the difference between responses to [6P4]CCL25 and WT CCL5. Proteins for which the 5P14-vs-WT response difference was significant (F test $P < 0.05$) but the 6P4-vs-WT response difference was not (F test $P > 0.25$) were selected. Proteins were additionally filtered for an increase (rather than a decrease) in proximity to CCR5 at both 3 min and 10 min post [5P14]CCL5 addition, relative to the basal state. Statistical significance calculations omitted the 10 min time point as potentially compromised by the poor quality of the [5P14]CCL5 10 min sample; however, the 10 min values were used in filtering.

Fig. S1.

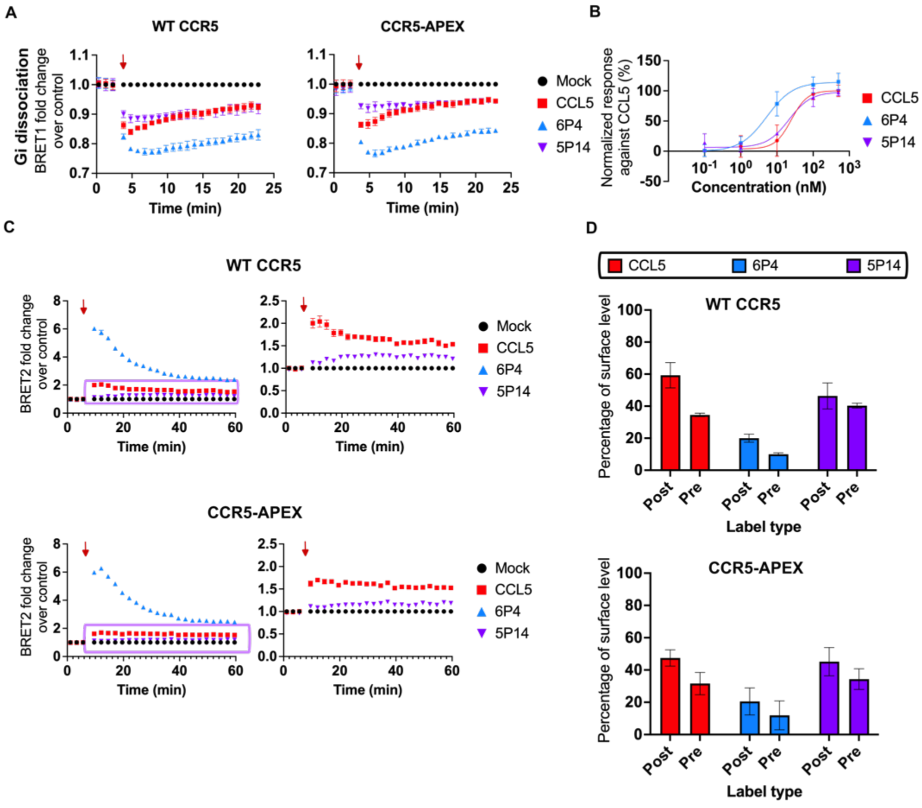


**Fig. S1. CCR5-APEX receptor exhibits wildtype-like signaling and trafficking.** (A) HEK293T cells expressing either WT CCR5 or APEX conjugated (CCR5-APEX) receptor and transfected with an IRES vector coding for Gαi-Nluc and Gβγ-cpVenus were stimulated with 100 nM CCL5, 6P4, or 5P14, respectively (indicated by red, blue and purple data points, respectively). The dissociation of the Gβγ from the Gα subunit results in a decrease in the BRET ratio upon ligand treatment (indicated by a red arrow) and is normalized against the CCL5 response (%). All data are expressed as the mean ± SE of n = 3 independent experiments. (B) G protein dissociation assays in (A) were performed using WT CCR5 expressing HEK293T cells with varying concentrations of CCL5, 6P4, or 5P14 (0, 1, 10, 100, 500 nM). The total G protein dissociation response was quantified and normalized to the response of CCL5. All data are expressed as the mean ± SE of n = 2 independent experiments. (C) Bystander BRET assays using luciferase-fused β-arrestin2 and GFP-fused CAAX in either WT CCR5 or CCR5-APEX expressing HEK293T cells showing β-arrestin2 recruitment to the plasma membrane (PM) after ligand stimulation (100nM CCL5, 6P4, or 5P14). Ligand addition is indicated by a red arrow and the normalized BRET signal is expressed as the fraction of the BRET ratio to that observed in the untreated control cells. The BRET signal traces from the CCL5 and 5P14 treated groups are indicated by a purple box and enlarged on the right. All data are expressed as the mean ± SE of n = 3 independent experiments. (D) Internalization of HEK293T cells stably expressing WT CCR5 or CCR5-APEX after ligand stimulation (100nM CCL5, 6P4, or 5P14) were assessed by pre-label flow cytometry (see Methods). Data are presented as fraction of surface receptor remaining compared to non-internalized control.

Fig.S2.

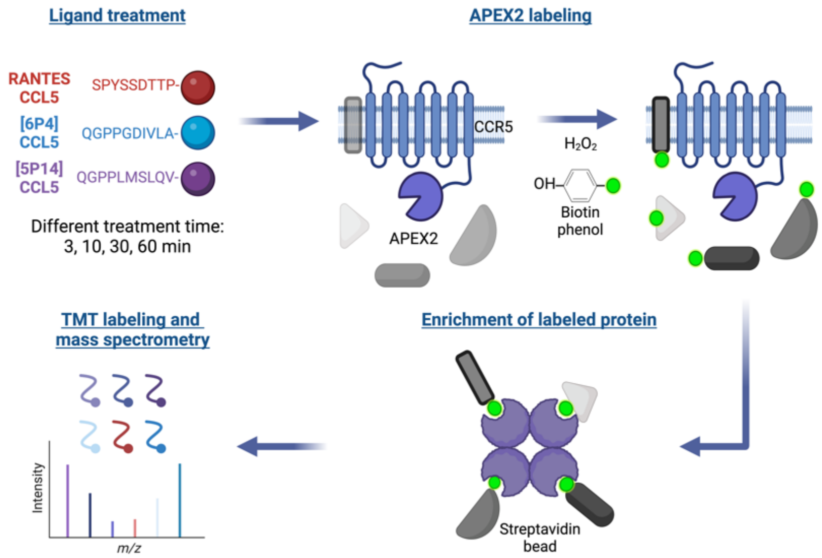


**Fig. S2. Experimental scheme of CCR5-APEX proximity labeling.** CCR5-APEX cells were pre-treated with biotin-phenol and then stimulated with 100nM of the corresponding CCL5 ligand over a time course of up to 60 minutes. At 3, 10, 30 and 60 minutes after stimulation, peroxide was added to initiate labeling and the reaction was quenched after 45 seconds. The labeled proteins were then extracted from cells and purified using streptavidin beads. All samples were then labeled with "tandem mass tag" fluorophore reagents and analyzed together in a single mass spectrometry experiment to ensure the best possible quantification, reproducibility and significance.

Fig. S3.

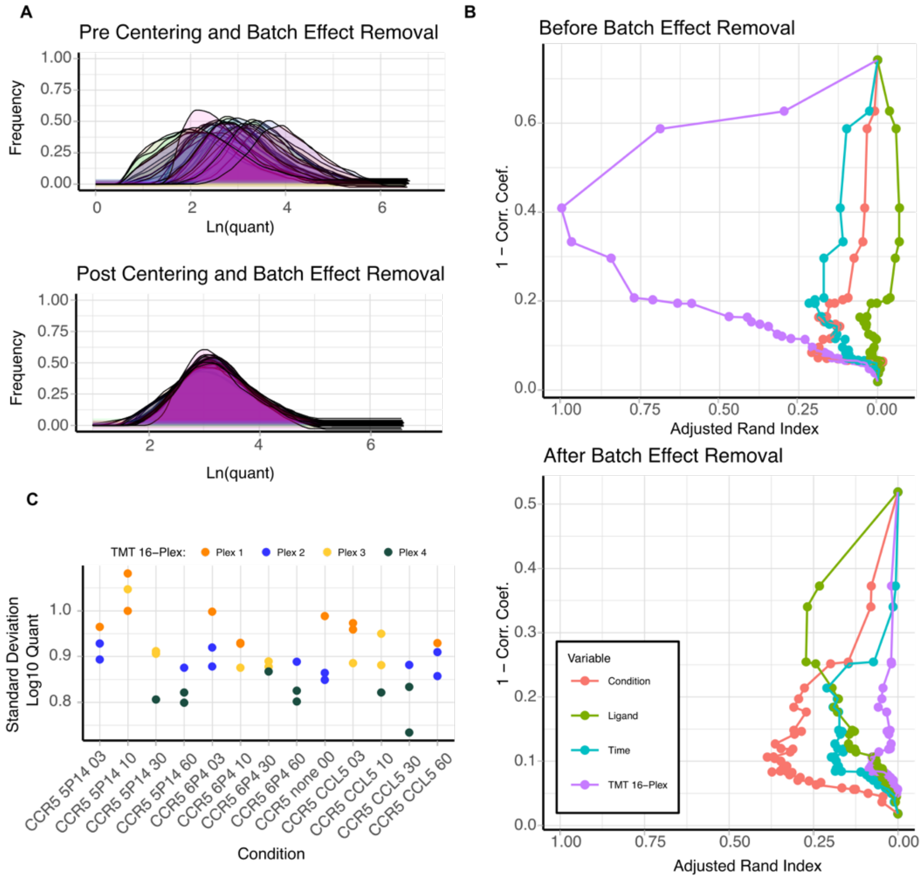


**Fig. S3. Data quality control for peptide-spectrum match quant analysis of CCR5-APEX proximity labeling proteomics.** (A) Peptide-spectrum match quant distribution for all samples before (upper panel) and after (lower panel) median centering and batch effect removal. (B) Adjusted Rand indices between the sample-to-sample correlation dendrogram cut at different heights and the experimental variables were compared before (upper panel) and after (lower panel) batch effect removal. (C) Standard deviations of peptide-spectrum match quant distributions for all ligand conditions. Each dot represents one sample. The numbers in condition names decode the time after ligand stimulation (00: no stimulation; 03: 3 min; 10: 10 min; 30: 30 min; 60: 60 min.) The different colors indicate data from different TMT 16-plex batches (4 batches total).

Fig. S4.

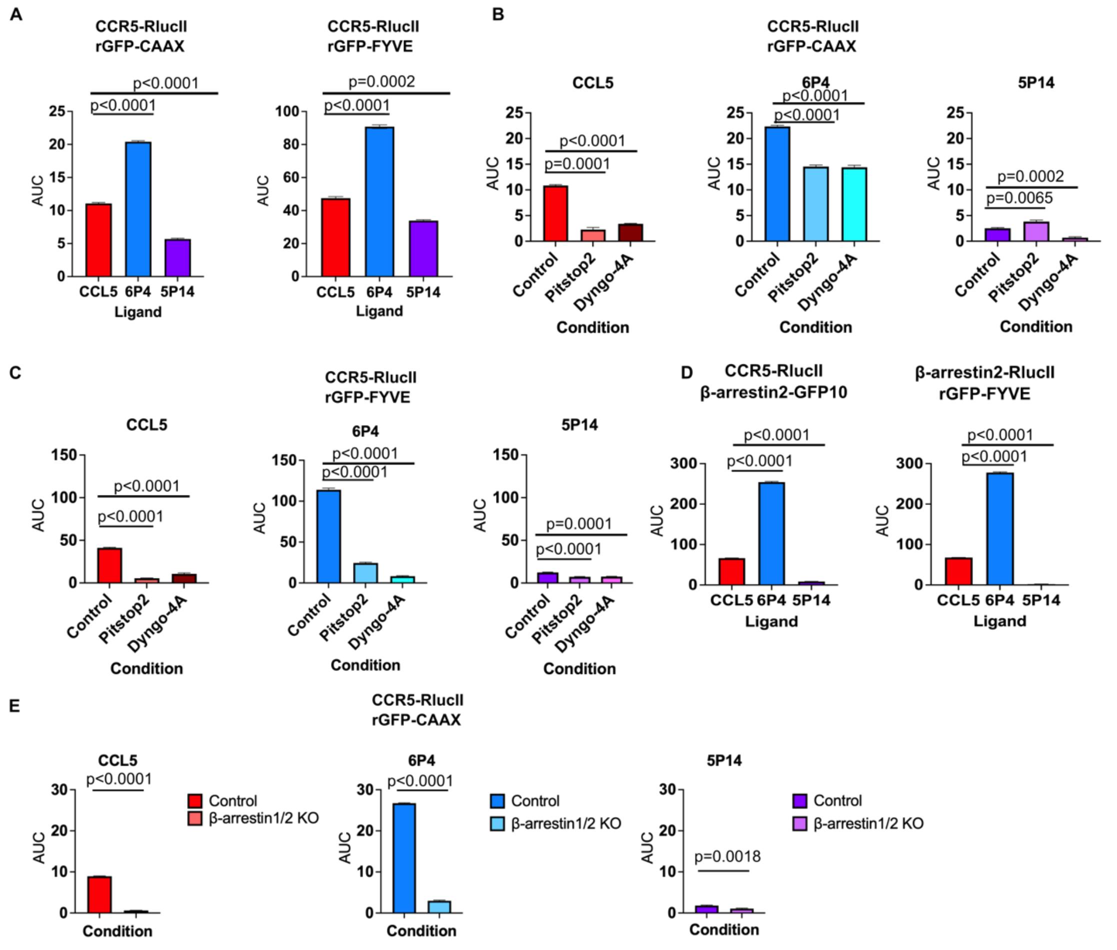


**Fig. S4. Area under curve quantification and statistical analysis of BRET assays in Fig. 2.** (A) Area under curve (AUC) analysis for bystander BRET between CCR5-RlucII and rGFP-CAAX or rGFP-FYVE (corresponding to Fig. 2A). (B-C) AUC analysis for bystander BRET between CCR5-RlucII and rGFP-CAAX (B) or rGFP-FYVE (C) with different clathrin-mediated endocytosis (CME) inhibitor treatments, Pitstop®2 and Dyngo-4a (corresponding to Fig. 2C-D). The same ligand treatment condition (100nM CCL5, 6P4, or 5P14) are plotted together to demonstrate the differences among inhibitor treatments. (D) AUC analysis for BRET between CCR5-RlucII and β-arrestin2-GFP10 or between RlucII-β-arrestin2 and rGFP-FYVE (corresponding to Fig. 2E and 2F). (E) AUC analysis for bystander BRET between CCR5-RlucII and rGFP-CAAX in control HEK293 cells or β-arrestin1/2 knockout (KO) HEK293 cells. The same ligand treatment condition (100nM CCL5, 6P4, or 5P14) are plotted together to demonstrate the differences between the control and the knockout cell line. All analyses were calculated with the baseline set to 1. All data are expressed as the mean ± SE of n = 3 independent experiments. P values were calculated using the unpaired t test with Welch's correction.

**Fig. S5.**

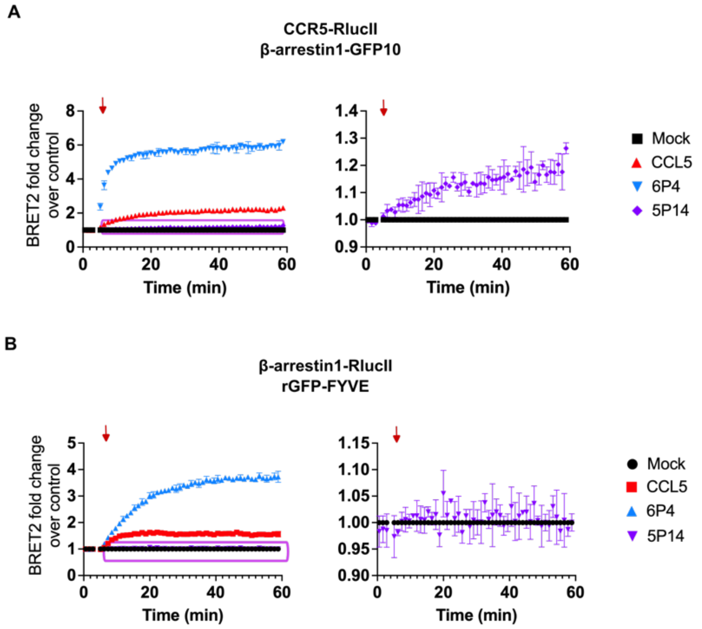


**Fig. S5. CCL5 variants distinctly induce β-arrestin1 recruitment to CCR5.** (A) Recruitment of β-arrestin1 to CCR5 using luciferase-fused CCR5-RlucII and β-arrestin1-GFP10 in HEK293T cells following treatment with 100nM CCL5, 6P4, or 5P14. (B) Recruitment of β-arrestin1 to endosomes using bystander BRET assays with β-arrestin1-RlucII and rGFP-FYVE in CCR5 expressing HEK293T cells after treatment with 100nM CCL5, 6P4, or 5P14). The ligand addition is indicated by a red arrow and the normalized BRET signal is expressed as fraction of the BRET ratio to that observed in the untreated control cells. The BRET signal traces from the 5P14 treated groups are indicated by a purple box and enlarged on the right. All data are expressed as the mean ± SE of n = 3 independent experiments.

Fig. S6.

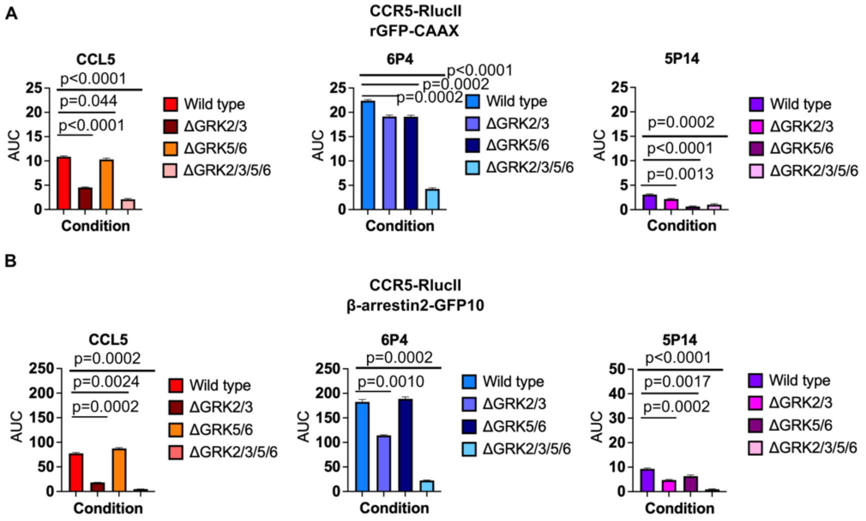


**Fig. S6. Area under curve quantification and statistical analysis of BRET assays in Fig. 3.** AUC analysis for BRET between CCR5-RlucII and rGFP-CAAX (A) or β-arrestin2-GFP10 (B) in parental HEK293A cells, ΔGRK2/3), ΔGRK5/6), or ΔGRK2/3/5/6 HEK293A cells. The same ligand treatment conditions (100nM CCL5, 6P4, or 5P14) are plotted together to demonstrate the differences between the control and the knockout cell lines. All analyses were calculated with the baseline at 1. All data are expressed as the mean ± SE of n = 3 independent experiments. P values were calculated using the unpaired t test with Welch's correction.

Fig. S7.

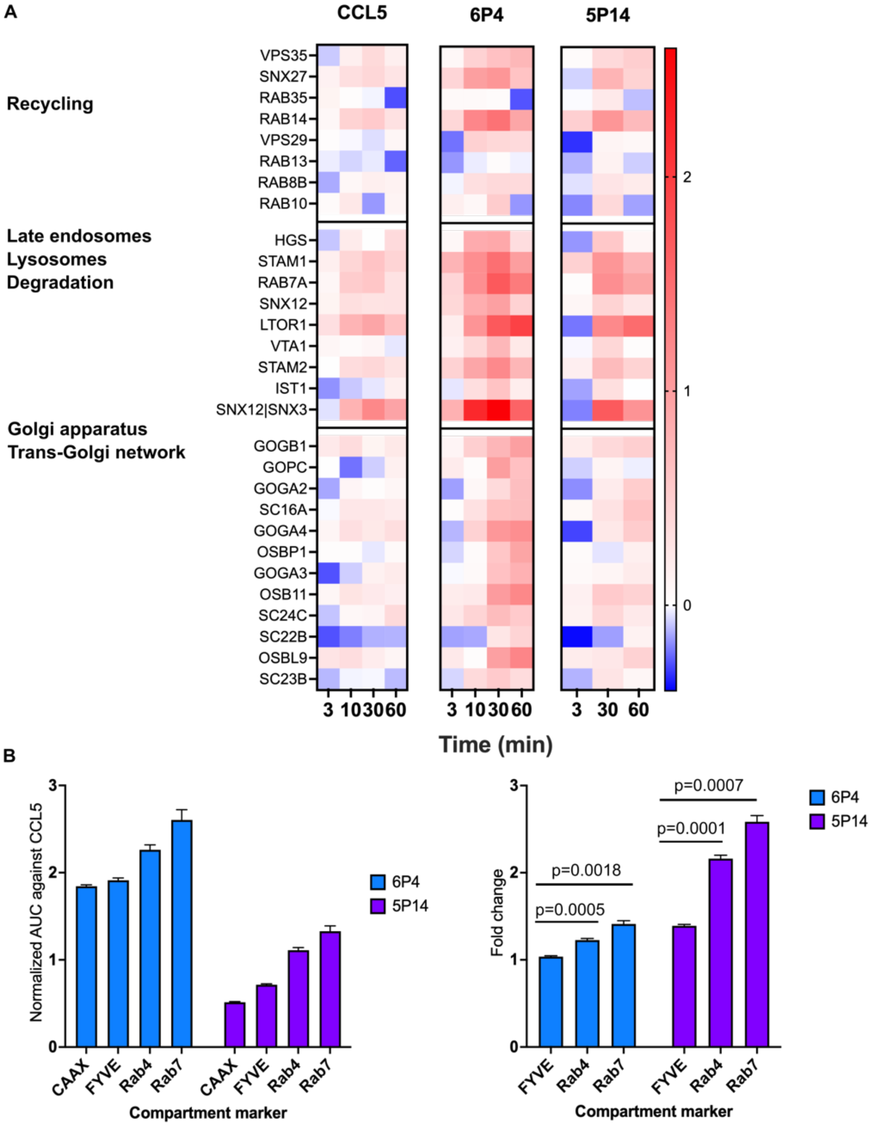


**Fig. S7. Comparisons of different trafficking pathways of CCR5 induced by different CCL5 variants.** (A) Heatmaps of time-dependent CCR5-APEX labeling of proteins associated with recycling, late endosomes, lysosomes, and endo-lysosomal degradation pathways and resident proteins in Golgi and Trans-Golgi network are plotted together for each CCL5 ligand. (B) Area under curve was calculated for bystander BRET assays between CCR5-RlucII and rGFP-CAAX, rGFP-FYVE, rGFP-Rab4 or rGFP-Rab7 with baselines set to 1. For each BRET assay, the AUC values of the 6P4- and 5P14-treated samples were normalized against the CCL5 condition and were then plotted on the left. The normalized AUC values in CCR5-RlucII and rGFP-FYVE, rGFP-Rab4 or rGFP-Rab7 BRET assay were then divided by the normalized AUC value of CCR5-RlucII and rGFP-CAAX to calculate the fold changes for both 6P4 and 5P14 conditions, which are plotted on the right. All data are expressed as the mean ± SE of n = 3 independent experiments. P values were calculated using the unpaired t test with Welch's correction.

**Fig. S8.**

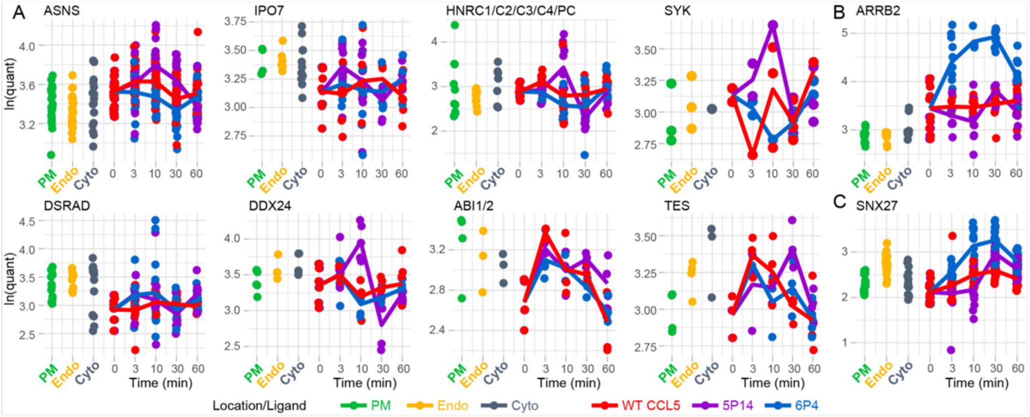


**Fig. S8. Response profiles of proteins whose proximity to CCR5 increase exclusively with [5P14]CCL5 stimulation and are distinct from WT CCL5 stimulation.** Protein proximities to CCR5 3 min and 10 min after the addition of [5P14]CCL5 were compared to the basal proximities. Filtered proteins (**Fig. 8A**) were selected using the significance of the difference in response to [5P14]CCL5 vs WT CCL5 ($P < 0.05$) and the significance of the difference in response to [6P4]CCL5 vs WT CCL5 ($P > 0.25$). (A) The resulting eight proteins (in addition to 1433ζ) are shown. (B) The proximity response of β-arrestin2 decreased, which is the opposite of the 1433ζ proximity response (**Fig 8B**), only with the [5P14]CCL5 ligand variant. (C) The increase in SNX27 proximity to CCR5 is delayed in response to [5P14]CCL5.

**Table S1. List of significantly varied proteins identified by two-way moderated ANOVA analysis with Benjamini-Hochberg (BH) adjustment.**

| Protein | -Log10 (Adjusted P-value) | Residual Degree of Freedom | F value |
|---|---|---|---|
| ARRB2 | 30.95337943 | 118 | 43.7260998 |
| SNX3 | 23.19322846 | 247 | 121.458957 |
| WASH4 | 22.16616134 | 77 | 25.3924852 |
| RAB7A | 21.10625739 | 327 | 81.9193279 |
| TFG | 20.59744077 | 75 | 23.0395194 |
| STX12 | 18.44381496 | 269 | 55.0038805 |
| RAB2A | 18.04636998 | 71 | 23.2674082 |
| PICAL | 18.04285714 | 97 | 24.3683827 |
| SCAM3 | 17.67002931 | 289 | 53.7248942 |
| ACTB | 17.15143098 | 75 | 17.5568228 |
| SNP23 | 17.03696371 | 52 | 28.6703305 |
| K1522 | 16.36571769 | 68 | 27.3431957 |
| OSB11 | 16.05378251 | 181 | 30.2133339 |
| P53 | 16.01846449 | 258 | 31.8271191 |
| ZFY16 | 15.71618317 | 192 | 15.8302198 |
| AAAT | 15.13902165 | 133 | 16.8323361 |
| STX7 | 15.0654394 | 155 | 26.0225664 |
| LTOR1 | 15.06464082 | 73 | 15.6002729 |
| WDR91 | 14.98859231 | 154 | 17.0703357 |
| STAM1 | 14.70400206 | 266 | 49.9005116 |
| KTN1 | 14.46181022 | 184 | 10.4176669 |
| TERA | 14.44747063 | 273 | 12.2327142 |
| GOPC | 14.1164024 | 195 | 12.3108494 |
| DAG1 | 14.0904143 | 158 | 39.6050164 |
| FLVC1 | 13.91759694 | 114 | 20.8781518 |
| FAS | 13.7404307 | 108 | 11.3814334 |
| TOLIP | 13.68888187 | 142 | 30.0779289 |
| CLH1 | 13.60448556 | 319 | 44.0799996 |
| RAC1 | 13.21626799 | 75 | 12.4339703 |
| TS101 | 13.21085635 | 230 | 8.53818822 |
| CLCA | 12.9370133 | 221 | 31.0515823 |
| IQGA1 | 12.72821037 | 243 | 33.5115977 |
| VTI1B | 12.67336016 | 125 | 12.0240599 |
| TBC8B | 12.44261233 | 30 | 23.7615824 |
| IST1 | 12.3797588 | 186 | 9.18692575 |

| | | | |
|---|---|---|---|
| SCRIB | 12.14325951 | 242 | 17.4946787 |
| LTV1 | 12.08660546 | 73 | 11.3782449 |
| TFR1 | 12.05534425 | 245 | 26.656705 |
| DSG2 | 12.00249887 | 328 | 15.5509145 |
| MARE1 | 11.97918162 | 193 | 13.5871184 |
| YTHD3 | 11.87431945 | 90 | 14.222771 |
| UBP10 | 11.69894764 | 287 | 20.6638711 |
| OSBP1 | 11.63836862 | 176 | 18.3280325 |
| ARRS | 11.57970328 | 20 | 38.9540318 |
| PVR | 11.53824847 | 32 | 20.6223316 |
| VIME | 11.52016464 | 108 | 9.12249352 |
| CTNB1 | 11.15147429 | 270 | 35.2779521 |
| OSBL9 | 11.1442549 | 102 | 17.7986133 |
| NUMB | 11.12021933 | 65 | 17.8527606 |
| FLNA | 10.95551904 | 104 | 19.7432181 |
| CNN2 | 10.7721964 | 187 | 9.85949058 |
| ITB1 | 10.64580573 | 109 | 15.7203702 |
| XRP2 | 10.61930515 | 67 | 16.742233 |
| P4K2A | 10.53192598 | 67 | 11.1665304 |
| IMA1 | 10.49347208 | 143 | 14.0300696 |
| FKB15 | 10.38320037 | 219 | 14.0198884 |
| RUVB1 | 10.31499416 | 117 | 9.93128635 |
| SCAM1 | 10.28024932 | 74 | 11.4894131 |
| SRC8 | 10.05045591 | 244 | 28.8615683 |
| STRAP | 9.911939698 | 189 | 16.4841784 |
| IQGA3 | 9.90702518 | 66 | 10.9976968 |
| ALBU | 9.878005625 | 97 | 8.11846644 |
| CBPD | 9.853574463 | 107 | 10.9031302 |
| AMOL1\|AMOT | 9.747287315 | 140 | 11.4444553 |
| ATP7B | 9.712051724 | 44 | 11.5848487 |
| AEDO | 9.638633791 | 74 | 8.74875682 |
| SNX12\|SNX3 | 9.528624362 | 20 | 25.8594205 |
| CTND1 | 9.438038161 | 241 | 16.0296694 |
| MYL6 | 9.405277902 | 38 | 12.2526753 |
| AT2B1\|AT2B4 | 9.320048713 | 42 | 12.083686 |
| PACN2 | 9.225009447 | 296 | 10.3998267 |
| FARP1 | 9.152216427 | 20 | 26.175132 |
| PTPRA | 9.07993659 | 44 | 21.4231187 |
| PTN23 | 9.041147044 | 256 | 23.0231365 |

| | | | |
|---|---|---|---|
| CLAP1\|CLAP2 | 8.976980348 | 88 | 7.64345383 |
| FKBP4 | 8.942002481 | 220 | 11.4970026 |
| KLRG2 | 8.913273201 | 68 | 11.749813 |
| GOGA3 | 8.904990363 | 147 | 16.9852221 |
| STAM2 | 8.861254507 | 149 | 20.0473676 |
| ESYT1 | 8.814648169 | 240 | 15.28624 |
| ATX2 | 8.745642199 | 188 | 11.7268045 |
| PALS2 | 8.720053941 | 88 | 10.287632 |
| HGS | 8.605795602 | 221 | 11.4824173 |
| CARM1 | 8.525112708 | 131 | 10.0466312 |
| CK5P2 | 8.39286671 | 97 | 7.64898806 |
| CP51A | 8.383042419 | 124 | 16.7230488 |
| MYH14 | 8.300440863 | 20 | 18.8823333 |
| AT1B3 | 8.272761161 | 98 | 8.12788779 |
| TLN2 | 8.19122599 | 30 | 11.969971 |
| ATG4B | 8.137732387 | 29 | 12.1578342 |
| ERBB2 | 8.06007744 | 32 | 11.985806 |
| ARFP1 | 8.020347268 | 18 | 18.0309621 |
| 1433Z | 7.945364793 | 99 | 6.57794716 |
| IF4G1 | 7.919106511 | 283 | 35.7410048 |
| SC23A | 7.906017932 | 227 | 25.2333964 |
| RALA\|RALB | 7.905255111 | 42 | 9.93796812 |
| VPS45 | 7.894806987 | 31 | 11.8165884 |
| RACK1 | 7.828125178 | 129 | 9.59088659 |
| FERM2 | 7.823554093 | 237 | 31.049048 |
| GOGA4 | 7.799620508 | 79 | 13.8335459 |
| SPTB2 | 7.783009828 | 197 | 7.75127197 |
| H33 | 7.766805989 | 71 | 7.13450409 |
| MARK2 | 7.751457244 | 43 | 22.4117177 |
| UTP6 | 7.67825435 | 96 | 6.91962288 |
| RRAS2 | 7.673855851 | 132 | 13.2743994 |
| SC24C | 7.668415035 | 197 | 10.4588904 |
| STX8 | 7.634596634 | 54 | 8.59406639 |
| RGAP1 | 7.626923382 | 165 | 8.02540278 |
| RAB14 | 7.596600911 | 165 | 26.5148459 |
| MRP | 7.591741991 | 71 | 6.98135121 |
| S12A2 | 7.59132171 | 205 | 12.592919 |
| VPS11 | 7.573743008 | 8 | 49.8925887 |
| VPS35 | 7.472841071 | 190 | 9.6395727 |

| | | | |
|---|---|---|---|
| GNAS1 | 7.450000449 | 220 | 13.5369724 |
| IRS4 | 7.44803604 | 320 | 14.0461111 |
| LAR4B | 7.405397605 | 172 | 7.78328692 |
| CKAP2 | 7.378588247 | 102 | 8.50447971 |
| ANXA7 | 7.31578738 | 344 | 15.7187476 |
| ZO2 | 7.26931031 | 258 | 5.33169926 |
| SMN | 7.259593196 | 177 | 15.2381712 |
| LRCH3 | 7.243575447 | 79 | 6.44828206 |
| P85A | 7.056422874 | 68 | 7.79756583 |
| AMOT | 7.024950354 | 292 | 8.23754284 |
| CATA | 7.001492671 | 66 | 6.61584435 |
| IQGA1\|IQGA2\|IQGA3 | 6.976893467 | 20 | 14.2138944 |
| YES | 6.945064227 | 213 | 14.44962 |
| SNX27 | 6.926168906 | 270 | 31.7064739 |
| RBP2 | 6.914876187 | 142 | 13.660317 |
| RAB9A\|RAB9B | 6.898138764 | 20 | 13.8305577 |
| GCP60 | 6.890516903 | 136 | 10.5274563 |
| ARF1\|ARF3\|ARF4\|ARF5\|ARF6\|ARL17 | 6.884645973 | 30 | 9.91475494 |
| ARL11 | 6.884645973 | 30 | 9.91475494 |
| IF2B1 | 6.853767049 | 253 | 10.4645574 |
| KCD12 | 6.801448344 | 297 | 7.04204818 |
| SNTB2 | 6.794512693 | 201 | 8.28085466 |
| NEB1 | 6.780341434 | 8 | 34.7261764 |
| WDR6 | 6.654442605 | 170 | 14.9089323 |
| H4 | 6.617929284 | 28 | 9.37854031 |
| ABI1 | 6.616990827 | 78 | 6.32748001 |
| NSA2 | 6.597997149 | 55 | 6.66987062 |
| SPTN1 | 6.522254808 | 237 | 10.2437235 |
| STX16 | 6.518436371 | 18 | 15.0646486 |
| OCLN | 6.513497503 | 76 | 10.7163629 |
| IQGA1\|IQGA2 | 6.492422295 | 30 | 8.7626117 |
| AKA12 | 6.476851879 | 190 | 19.9771525 |
| RFIP1 | 6.474741731 | 132 | 15.1339353 |
| TXLNG | 6.464625713 | 233 | 9.32962654 |
| NISCH | 6.448902421 | 147 | 11.0920736 |
| WNK1\|WNK2\|WNK4 | 6.426953952 | 75 | 5.87361884 |
| CTNA1 | 6.390126634 | 202 | 25.1370398 |
| MYPT1 | 6.352758242 | 198 | 4.84441915 |
| PUM1\|PUM2 | 6.307758572 | 75 | 6.42769273 |

| | | | |
|---|---|---|---|
| RTN3 | 6.271776492 | 75 | 6.30914199 |
| COBL1 | 6.244340093 | 45 | 6.82921564 |
| FYN | 6.236372182 | 88 | 7.96507597 |
| CDV3 | 6.226443115 | 143 | 14.0545373 |
| LSR | 6.222855499 | 75 | 15.3794269 |
| ANX11 | 6.216452389 | 288 | 12.5991493 |
| CSKP | 6.20137113 | 169 | 14.5169404 |
| FARP2 | 6.106318924 | 30 | 8.63683301 |
| RAC1\|RAC2\|RAC3 | 6.07656266 | 63 | 6.25698314 |
| ZFYV9 | 6.056623943 | 147 | 10.4896192 |
| REPS1 | 6.037570375 | 239 | 9.06719013 |
| DCTN2 | 5.992359309 | 252 | 8.72534027 |
| PLAK | 5.980507505 | 292 | 11.6671596 |
| LMAN2 | 5.969813517 | 30 | 7.94187294 |
| AT2B1 | 5.957553768 | 117 | 10.7681744 |
| RAB1A | 5.914204363 | 129 | 6.13193201 |
| NECT2 | 5.849830199 | 68 | 5.8765114 |
| RHG17 | 5.828556772 | 18 | 11.0510041 |
| GOGB1 | 5.822425786 | 194 | 15.7967885 |
| ABI1\|ABI2 | 5.822271386 | 42 | 6.96911216 |
| GNA11\|GNAQ | 5.806011044 | 20 | 10.9173147 |
| MD2L1 | 5.778616757 | 20 | 10.8657587 |
| NEK9 | 5.76391672 | 181 | 4.98488524 |
| CROCC | 5.746562298 | 30 | 8.10695382 |
| AT11C | 5.746213896 | 173 | 11.526195 |
| VTA1 | 5.736921907 | 55 | 10.6623765 |
| KIF23 | 5.710124055 | 184 | 4.52148583 |
| TNPO1\|TNPO2 | 5.687542715 | 112 | 6.87771307 |
| BI2L1 | 5.687429272 | 187 | 17.2547283 |
| RAGP1 | 5.664188562 | 269 | 17.4174259 |
| RASN | 5.653557702 | 97 | 7.11649656 |
| RHEB | 5.625759335 | 19 | 10.3751403 |
| NOTC2 | 5.533982845 | 193 | 17.6600266 |
| EPN4 | 5.513204936 | 126 | 11.8960537 |
| SNX12 | 5.508767495 | 77 | 6.75541379 |
| VPS29 | 5.450867552 | 99 | 7.49986064 |
| VAPA | 5.44488926 | 51 | 5.69568656 |
| HIP1R | 5.441042893 | 202 | 7.23727103 |
| TGFA1 | 5.417423256 | 8 | 21.9341374 |

| | | | |
|---|---|---|---|
| WASF2 | 5.410506261 | 214 | 10.2025886 |
| CADH2 | 5.409355975 | 212 | 13.3352265 |
| SMAD5 | 5.405804512 | 53 | 5.58435598 |
| TXLNA | 5.394225131 | 94 | 7.91304748 |
| SNX9 | 5.384352419 | 81 | 10.0265759 |
| P20D2 | 5.35584084 | 41 | 6.73788291 |
| SC24B | 5.343919387 | 213 | 9.72932276 |
| SC5A6 | 5.333452467 | 18 | 9.62112183 |
| PROF1 | 5.293633362 | 342 | 13.7800658 |
| RAB5C | 5.25462844 | 236 | 10.7535447 |
| NIBA2 | 5.253435692 | 64 | 5.1477261 |
| PCDH7 | 5.219738483 | 142 | 16.9570342 |
| PHB1 | 5.21435328 | 312 | 12.6003416 |
| LRRC1\|SCRIB | 5.203041688 | 42 | 6.42673893 |
| MAK16 | 5.202392467 | 38 | 6.08265138 |
| AIMP2 | 5.200559276 | 87 | 6.38563931 |
| FA98A | 5.160741218 | 53 | 5.35151817 |
| TB10B | 5.159691782 | 104 | 7.43438564 |
| SRPRB | 5.127353719 | 96 | 5.98143063 |
| SMAP2 | 5.120972074 | 51 | 5.45971974 |
| CP110 | 5.104743889 | 8 | 19.7597887 |
| PLCB3 | 5.098699813 | 87 | 5.56533186 |
| AP2B1 | 5.08130644 | 122 | 7.58520848 |
| EIF3J | 5.079181349 | 112 | 5.4894998 |
| ABCE1 | 5.065904251 | 229 | 15.1710168 |
| SMAD2\|SMAD3\|SMAD9 | 5.052943096 | 30 | 6.5477884 |
| ALDOC | 5.037370768 | 60 | 5.05238831 |
| GAB1 | 5.037257999 | 64 | 7.78503978 |
| OBI1 | 5.009893637 | 90 | 4.5718987 |
| TGO1 | 5.005835541 | 168 | 7.05130201 |
| CLIC4 | 5.004322322 | 8 | 17.2761329 |
| DLG1\|DLG2 | 4.980356605 | 30 | 6.44240533 |
| DCP1B | 4.952516335 | 28 | 6.66944347 |
| RALA | 4.949722409 | 44 | 5.47763705 |
| ZO1 | 4.934252421 | 307 | 8.41285282 |
| ARF5 | 4.930309251 | 30 | 6.37063871 |
| STXB3 | 4.919854822 | 157 | 7.78727041 |
| ROBO1\|ROBO2 | 4.907181139 | 8 | 16.7571622 |
| EGFR\|ERBB2 | 4.905310973 | 20 | 7.82594009 |

| | | | |
|---|---|---|---|
| REV1 | 4.905310973 | 18 | 8.34184403 |
| TMF1 | 4.90496294 | 54 | 5.08029017 |
| DLDH | 4.891220407 | 18 | 8.30413914 |
| SYAP1 | 4.891093415 | 128 | 9.67600051 |
| KC1G1\|KC1G2\|KC1G3 | 4.864837561 | 19 | 8.60094381 |
| ADAM9 | 4.860355959 | 40 | 5.56446143 |
| CLCB | 4.852378521 | 86 | 6.05106201 |
| NAF1 | 4.821985946 | 30 | 6.21351942 |
| EMD | 4.787968047 | 183 | 7.74398936 |
| PEF1 | 4.762329128 | 200 | 5.96740884 |
| ARF4 | 4.745765788 | 30 | 6.11363473 |
| EPS15 | 4.741523636 | 132 | 11.3979213 |
| OSBL8 | 4.733191001 | 165 | 5.91412502 |
| RAB13 | 4.704968436 | 134 | 6.95990714 |
| PUR8 | 4.689284746 | 130 | 4.42095903 |
| RHG12 | 4.653520319 | 32 | 6.2204712 |
| GOGA2 | 4.651570017 | 109 | 9.578817 |
| ARL8B | 4.63533379 | 19 | 7.73207119 |
| ZDHC5 | 4.628077748 | 76 | 5.59328891 |
| KLC1 | 4.618856463 | 111 | 4.95690209 |
| ATX10 | 4.612884864 | 163 | 6.90511335 |
| EDC3 | 4.600924848 | 135 | 4.21867156 |
| DESP | 4.59459931 | 276 | 5.74896789 |
| KIF14 | 4.558056888 | 180 | 3.87108638 |
| ACTN1 | 4.522843183 | 184 | 9.14353243 |
| PURA2 | 4.477353784 | 158 | 9.40256178 |
| BAG3 | 4.474943514 | 154 | 4.3660543 |
| DDX56 | 4.455081276 | 175 | 3.82359245 |
| WDR44 | 4.44042069 | 75 | 4.31754521 |
| KCC2A\|KCC2B\|KCC2D\|KCC2G | 4.435480273 | 30 | 5.7041869 |
| RPAB3 | 4.389825171 | 8 | 14.4887428 |
| PCAT1 | 4.362587032 | 62 | 4.43959364 |
| DHB12 | 4.355014719 | 98 | 4.04828104 |
| FLOT1 | 4.341404031 | 32 | 5.66010644 |
| UBL5 | 4.339210855 | 30 | 5.66712824 |
| RABE1 | 4.334017312 | 145 | 8.19668965 |
| GLGB | 4.292717184 | 29 | 5.58997502 |
| DDX52 | 4.28934026 | 115 | 6.18013636 |
| SPD2B | 4.283858679 | 136 | 4.25269191 |

| | | | |
|---|---|---|---|
| STK3\|STK4 | 4.283366926 | 7 | 13.0451502 |
| WDR81 | 4.281288746 | 65 | 4.55029298 |
| RBMS1\|RBMS2 | 4.27852855 | 19 | 6.80497785 |
| MCMBP | 4.276740486 | 111 | 4.18991897 |
| VATB2 | 4.262036026 | 99 | 5.00771984 |
| CITE2 | 4.251357116 | 20 | 6.68163069 |
| EAA1 | 4.231410443 | 30 | 6.58641113 |
| HIP1 | 4.216480251 | 83 | 6.60483877 |
| RM14 | 4.213705971 | 42 | 4.79987902 |
| SC23A\|SC23B | 4.204219072 | 40 | 4.8654533 |
| KCC2D | 4.190578941 | 57 | 4.38466724 |
| FYN\|SRC\|YES | 4.189252019 | 28 | 5.71994327 |
| GDIB | 4.189050141 | 174 | 6.8867945 |
| MAGD1 | 4.15002869 | 232 | 4.35140325 |
| DREB | 4.138738412 | 318 | 10.6690663 |
| KLC1\|KLC2 | 4.134731791 | 20 | 6.59560913 |
| ODP2 | 4.131442088 | 102 | 6.80885261 |
| SDK2 | 4.12978177 | 20 | 6.8105128 |
| ARF3 | 4.110697215 | 117 | 3.95599695 |
| PFD3 | 4.074889657 | 161 | 5.13553059 |
| BAIP2 | 4.070043131 | 202 | 7.90822206 |
| SNAG | 4.053844722 | 80 | 3.97557348 |
| ANKL2 | 4.037047977 | 134 | 6.10294189 |
| TNR6A | 4.037003389 | 77 | 6.56736477 |
| RAB35 | 4.034522658 | 225 | 14.0533351 |
| VATA | 4.032034821 | 153 | 5.02192027 |
| KCRB | 4.026655034 | 281 | 3.45196515 |
| UBP8 | 4.009022188 | 42 | 4.59439575 |
| S39AA | 4.002956017 | 19 | 7.27063597 |
| CD81 | 3.990409845 | 18 | 7.04991095 |
| BCCIP | 3.981821293 | 30 | 6.35612806 |
| SC24A | 3.976494447 | 213 | 14.8571855 |
| DOC11 | 3.964697993 | 119 | 4.47065561 |
| SUMO1 | 3.962437665 | 36 | 5.15377805 |
| COPA | 3.962264482 | 285 | 3.53059502 |
| GNAI3 | 3.960867873 | 100 | 5.02504685 |
| ODPB | 3.96053124 | 115 | 5.36109489 |
| UBP14 | 3.949780665 | 32 | 4.97935166 |
| SYDM | 3.94707694 | 30 | 5.0732514 |

| | | | |
|---|---|---|---|
| ZN207 | 3.931073562 | 20 | 6.00913951 |
| RB11A | 3.883538868 | 182 | 3.47426171 |
| STIP1 | 3.878454089 | 213 | 5.97492895 |
| HACL2 | 3.868634439 | 39 | 4.5509938 |
| TRI25 | 3.867467823 | 99 | 4.49813562 |
| GRIN1 | 3.866295967 | 207 | 9.2975373 |
| UBN2 | 3.865920556 | 20 | 6.41823461 |
| FYN\|LCK\|SRC\|YES | 3.856061153 | 19 | 6.47112711 |
| SHLB1 | 3.850610033 | 193 | 5.47252848 |
| HMOX2 | 3.839035036 | 97 | 10.1767146 |
| DRG2 | 3.828991833 | 54 | 4.91531816 |
| TSN | 3.82280711 | 65 | 3.95545571 |
| RNF11 | 3.822776425 | 42 | 4.55745085 |
| PKHA5 | 3.794744156 | 213 | 5.96401287 |
| AKAP8 | 3.787415517 | 62 | 3.96466549 |
| ERBIN | 3.766028808 | 184 | 25.4548468 |
| THOP1 | 3.765523305 | 65 | 3.90629666 |
| DYN2 | 3.75775955 | 224 | 4.99945651 |
| NDUA4 | 3.7539034 | 65 | 3.89637978 |
| 3MG | 3.746714619 | 29 | 4.87647443 |
| ANK3 | 3.744236696 | 211 | 5.89812828 |
| MP2K1\|MP2K2 | 3.743181274 | 44 | 4.26238012 |
| TNPO1 | 3.740718004 | 110 | 4.03104487 |
| LCAP | 3.731318093 | 8 | 9.39047214 |
| KLC2 | 3.727819157 | 8 | 10.3760658 |
| BUB1B | 3.727782785 | 222 | 4.22851335 |
| CNOT2 | 3.717991783 | 42 | 4.41939893 |
| COPD | 3.717971623 | 256 | 6.49382749 |
| SYNC | 3.707688631 | 32 | 4.6623737 |
| RT05 | 3.668023448 | 38 | 4.65648694 |
| CDC42 | 3.6632963 | 147 | 5.08184053 |
| RB33B | 3.644466543 | 42 | 4.21934601 |
| FLOT2 | 3.639624176 | 67 | 5.45297749 |
| SL7A1 | 3.636253248 | 44 | 4.15679312 |
| CSK22 | 3.636253248 | 89 | 3.59800593 |
| NCKP1 | 3.613370471 | 287 | 7.93642151 |
| YTHD1 | 3.610789406 | 20 | 5.87543817 |
| UHRF1 | 3.607334567 | 30 | 4.64009153 |
| SC23B | 3.606216162 | 102 | 6.97091263 |

| | | | |
|---|---|---|---|
| GNA11 | 3.57972298 | 59 | 4.02670406 |
| SYYM | 3.576725508 | 6 | 11.5487434 |
| CAZA1 | 3.544306373 | 105 | 4.43757792 |
| RENT2 | 3.544062692 | 19 | 5.53611739 |
| M4K4\|MINK1\|TNIK | 3.542866041 | 123 | 4.67981712 |
| AKT2 | 3.536501561 | 44 | 4.06204209 |
| HSPB1 | 3.527531027 | 52 | 4.24471225 |
| LYN | 3.518895507 | 215 | 8.63570124 |
| AP2A1\|AP2A2 | 3.518763103 | 84 | 5.7567736 |
| DHX36 | 3.513730362 | 229 | 5.31269807 |
| DOCK1 | 3.507822617 | 8 | 9.77066099 |
| K1671 | 3.499547406 | 43 | 4.05012563 |
| PKN3 | 3.473403951 | 20 | 5.69778488 |
| LAP2B | 3.468511038 | 77 | 3.55751903 |
| ABCD3\|GEN | 3.460377523 | 20 | 5.52860552 |
| SHIP1 | 3.460377523 | 20 | 5.52860552 |
| HHIP | 3.460377523 | 20 | 5.52860552 |
| PCNT | 3.455187151 | 110 | 6.48286312 |
| MRCKA\|MRCKB | 3.44426947 | 39 | 4.12360816 |
| SH3G1 | 3.442084456 | 158 | 6.1805263 |
| SC31A | 3.437912988 | 164 | 10.557369 |
| DLG1\|DLG2\|DLG4 | 3.430966739 | 19 | 5.27760112 |
| CHIP | 3.428975084 | 42 | 4.23148203 |
| VRK2 | 3.413183718 | 29 | 4.69801119 |
| PUM1 | 3.401832846 | 177 | 3.19030645 |
| DLG1 | 3.380873433 | 251 | 19.6924447 |
| PSMD3 | 3.369005743 | 270 | 4.44366411 |
| SNX6 | 3.368352714 | 88 | 4.96468434 |
| CHAC2 | 3.366735359 | 30 | 4.53373699 |
| LIPB1 | 3.36133455 | 29 | 4.65046668 |
| CKAP5 | 3.357580903 | 219 | 6.96738032 |
| TCPB | 3.350867339 | 27 | 4.49465864 |
| PTN14 | 3.346654805 | 19 | 5.62724251 |
| NVL | 3.338809031 | 30 | 4.4889465 |
| SHLB1\|SHLB2 | 3.335842337 | 20 | 5.01319939 |
| GRAP1 | 3.335064067 | 32 | 4.22979775 |
| ATG3 | 3.333600411 | 206 | 6.95014242 |
| GRDN | 3.327287274 | 115 | 3.27307829 |
| DDI2 | 3.321651783 | 40 | 4.28359896 |

| | | | |
|---|---|---|---|
| TIAR | 3.314439495 | 29 | 4.33812298 |
| IQGA1\|IQGA3 | 3.306671857 | 8 | 8.95399418 |
| SRPK1 | 3.297008939 | 108 | 4.33382447 |
| ANM3 | 3.289410742 | 205 | 4.28506696 |
| P85B | 3.284897426 | 90 | 3.74137765 |
| AKAP9 | 3.282414046 | 179 | 16.6678154 |
| AAAS | 3.27399428 | 153 | 6.30524643 |
| WNK1 | 3.273627612 | 168 | 4.92553265 |
| KITH | 3.262241253 | 53 | 3.64467153 |
| PSME3 | 3.261480679 | 51 | 3.67350969 |
| CFA47 | 3.253039995 | 20 | 4.87972085 |
| MEAK7 | 3.243321333 | 8 | 8.54909731 |
| KC1E | 3.236771559 | 43 | 3.79823359 |
| TTC37 | 3.232136626 | 7 | 8.57275473 |
| ARF3\|ARF5 | 3.232046021 | 30 | 4.19462234 |
| XIAP | 3.225697367 | 39 | 6.25742382 |
| PHLB1 | 3.214779886 | 72 | 4.33001097 |
| AMOL1 | 3.213929159 | 129 | 3.15819162 |
| TTC1 | 3.213929159 | 32 | 4.08951278 |
| PARP1 | 3.213211411 | 256 | 6.15024906 |
| ASPP2 | 3.2106903 | 210 | 5.75224471 |
| IQGA2 | 3.206714515 | 170 | 11.7350269 |
| DIC | 3.202661404 | 52 | 3.73119838 |
| TANC1 | 3.200077522 | 8 | 7.4241892 |
| AAPK1\|AAPK2 | 3.178646838 | 30 | 4.1302021 |
| ARFG3 | 3.177801507 | 148 | 5.03308567 |
| TSC1 | 3.159028579 | 20 | 4.89685054 |
| PATL1 | 3.152605578 | 53 | 5.10764932 |
| PKP2 | 3.148842343 | 169 | 4.4173954 |
| HYCCI | 3.147581363 | 20 | 5.14410528 |
| Z3H7A | 3.146257922 | 19 | 4.91976717 |
| DMD | 3.112467889 | 44 | 3.89767141 |
| ROBO1 | 3.105403578 | 32 | 4.26648012 |
| AKP13 | 3.095513229 | 44 | 6.27828614 |
| BAG4 | 3.075483906 | 64 | 3.36643681 |
| MGST3 | 3.064656418 | 39 | 6.65043307 |
| S4A7 | 3.0541605 | 30 | 5.27429263 |
| RN214 | 3.038038838 | 17 | 4.8576722 |
| MYH9 | 3.030241017 | 181 | 7.47916505 |

| | | | |
|---|---|---|---|
| PTPRF | 3.024939368 | 183 | 5.88341046 |
| AGFG1 | 3.002561424 | 41 | 3.61995856 |
| HAUS4 | 3.001795722 | 8 | 7.49377377 |

**Table S2. Representative time-course clusters of CCL5 (red), 6P4 (blue), 5P14 (purple) treated CCR5-APEX labeled proteins.**

| Cluster Profiles 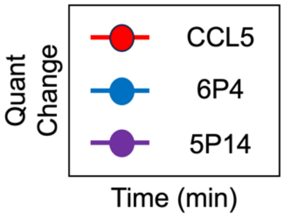 | Enriched Gene Ontology CC Terms | Representative proteins |
|---|---|---|
| 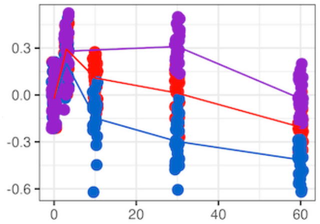 | Plasma membrane<br>Cell-cell junction<br>Extracellular exosome | CTND1, WASF2, MRP, AT1A1 |
| 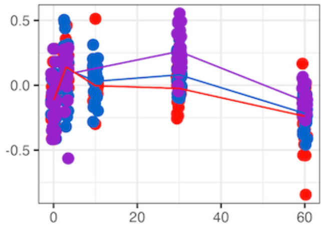 | Plasma membrane<br>Cell-cell junction<br>Actin-based cell projection | ITB1, DMD, EPS8, EM55, ZO1, IRS4, DSG2, LYN, PI4KA |
| 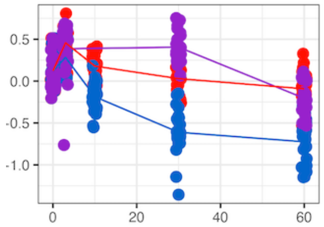 | Plasma membrane<br>Cell-cell junction | E41L3, AFAD, DLG1, XRP2, AT2B4, PHLB2, S39A6 |
| 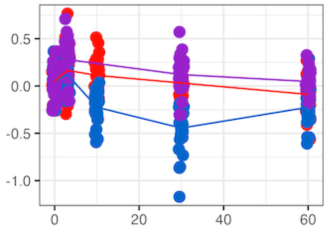 | Plasma membrane<br>Cell-cell junction<br>Cytoskeleton | DMD, GNAS1, GNA11, IQGA3, GRIN1, AKT2, EM55, PALM2, CTNL1, GNAI3, S12A2, MYO1D, YES, EPB41, CROCC, WASF2, AAAT, CLIC1, FLNB, STXB3, E41L5 |
| 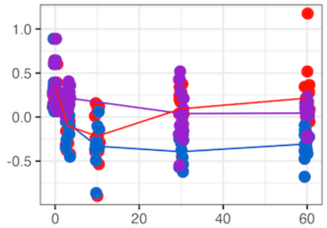 | Plasma membrane<br>Cell-cell junction<br>Myosin complex | CXAR, ZDHC5, NECT2, TB10B, BORG5, RALA, ROBO1, EFNB1, OST48, XRN1, ITA5, PHLB2, NED4L, ANK2, PP1B, YTDC1, MPP7, MYH10, C2CD5, S39AA, PALM2, VANG2, NEB2 |

| | | |
|---|---|---|
| 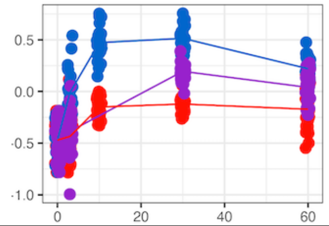  | Early endosome | PTN23, STAM2, RAB5C, CFA47, YTHD1/2/3 |
| 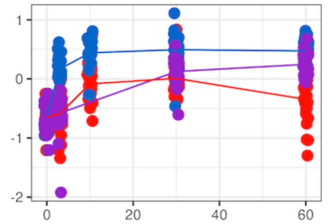  | Endosome<br>Clathrin coat | CLH1, VPS45, EPN4, RNF11, TGFA1, RMC1 |
| 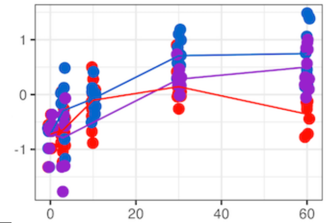  | Late endosome<br>Lysosome | TGFA1, LTOR1, AKTS1, SYYM, RMC1, RAB9A/B, LRBA/NBEA, RBM3, ARL8B, HNRC1, PAPS2, STX16, TTC37, P4K2A, HSP7E |
| 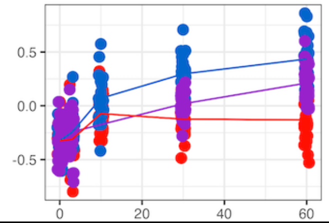  | Lysosome<br>Perinuclear region | AKAP9, SC16A, VPS35, RMC1, TOM1, Z3H7A, VPP1, WDR81, KTN1, MFR1L, ATX2, GCC2, VATA, GOGB1 |
| 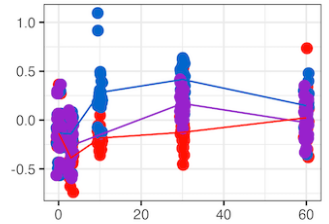  | Endoplasmic reticulum exit site<br>Intracellular vesicle | SC23A, EEA1, IST1, RB33B, RAB15, RAB10, WINK2, SC23B, RBMS1/2, TIAR, CE290, YTHD1, SNX24, SC24B, RAB1A, VPS16, SC24C, GRAN, HAUS4, F168A, GRAP1 |
| 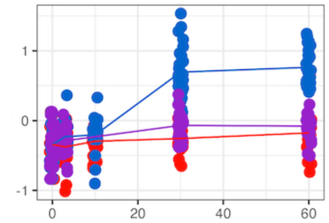  | Late endosome<br>Golgi apparatus<br>Trans-Golgi network | LAR4B/P4, RMC1, OSB11, ATP7B, OSBL9 |
| 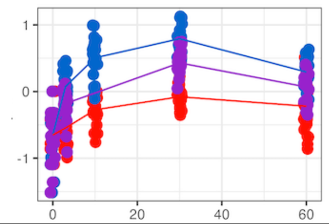  | Endosome<br>Golgi membrane | STAM1, RAB14, VPS45, ANX11/A7, TGFA1, LMAN2, CFA47, TBC8B |

| | | |
|---|---|---|
| 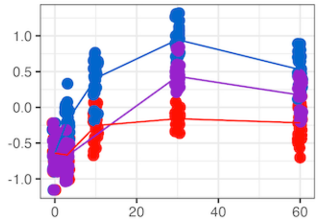  | Endosome<br>Golgi apparatus<br>Trans-Golgi network | STX12, TGFA1, SCAM3, RAB7A |
| 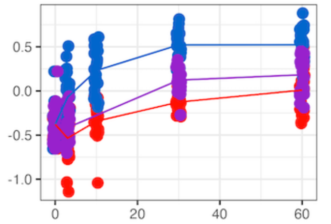  | Golgi apparatus<br>Intracellular vesicle | CLCA, YIPF5, KATL2, FKB15, TOLIP, ATP7B, LAR4B/LARP4 |

**References**


1. A. Campeau *et al.*, Multi-omics of human plasma reveals molecular features of dysregulated inflammation and accelerated aging in schizophrenia. *Molecular Psychiatry* **27**, 1217-1225 (2022).
2. M. E. Ritchie *et al.*, limma powers differential expression analyses for RNA-sequencing and microarray studies. *Nucleic Acids Res* **43**, e47 (2015).
3. G. K. Smyth, "limma: Linear Models for Microarray Data" in Bioinformatics and Computational Biology Solutions Using R and Bioconductor, R. Gentleman, V. J. Carey, W. Huber, R. A. Irizarry, S. Dudoit, Eds. (Springer New York, New York, NY, 2005), 10.1007/0-387-29362-0_23, pp. 397-420.
4. D. P. Nusinow, S. P. Gygi, A Guide to the Quantitative Proteomic Profiles of the Cancer Cell Line Encyclopedia. *Biorxiv* 10.1101/2020.02.03.932384, 2020.2002.2003.932384 (2020).
5. D. Mellacheruvu *et al.*, The CRAPome: a contaminant repository for affinity purification–mass spectrometry data. *Nature Methods* **10**, 730-736 (2013).
6. H. W. L. Koh *et al.*, EBprot: Statistical analysis of labeling-based quantitative proteomics data. *Proteomics* **15**, 2580-2591 (2015).
7. H. W. L. Koh, Y. Zhang, C. Vogel, H. Choi, EBprotV2: A Perseus Plugin for Differential Protein Abundance Analysis of Labeling-Based Quantitative Proteomics Data. *Journal of Proteome Research* **18**, 748-752 (2018).
8. T. Wu *et al.*, clusterProfiler 4.0: A universal enrichment tool for interpreting omics data. *Innovation (Camb)* **2**, 100141 (2021).
9. H. Motulsky, A. Christopoulos, *Fitting models to biological data using linear and nonlinear regression: a practical guide to curve fitting* (Oxford University Press, New York, 2004).
